# A systematic comparison of computational methods for expression forecasting

**DOI:** 10.1101/2023.07.28.551039

**Authors:** Eric Kernfeld, Yunxiao Yang, Joshua S. Weinstock, Alexis Battle, Patrick Cahan

## Abstract

Expression forecasting methods use machine learning models to predict how a cell will alter its transcriptome upon perturbation. Such methods are enticing because they promise to answer pressing questions in fields ranging from developmental genetics to cell fate engineering and because they are a fast, cheap, and accessible complement to the corresponding experiments. However, the absolute and relative accuracy of these methods is poorly characterized, limiting their informed use, their improvement, and the interpretation of their predictions. To address these issues, we created a benchmarking platform that combines a panel of 11 large-scale perturbation datasets with an expression forecasting software engine that encompasses or interfaces to a wide variety of methods. We used our platform to systematically assess methods, parameters, and sources of auxiliary data, finding that performance strongly depends on the choice of metric, and especially for simple metrics like mean squared error, it is uncommon for expression forecasting methods to out-perform simple baselines. Our platform will serve as a resource to improve methods and to identify contexts in which expression forecasting can succeed.

## Introduction

Genetic screening is of fundamental importance to basic biology, and in drug development, it roughly doubles the chance that a preclinical finding will survive translation (King, Davis, & Degner, 2019; Minikel, Painter, Dong, & Nelson, 2024; Nelson et al., 2015). Fueled by ATAC-seq (Buenrostro, Giresi, Zaba, Chang, & Greenleaf, 2013), single-cell RNA-seq (Ziegenhain et al., 2017), and related perturbation assays (Adamson et al., 2016; Datlinger et al., 2017; Dixit et al., 2016; Jaitin et al., 2016), a raft of recent computational methods now offer *expression forecasting*: prediction of genetic perturbation effects on the transcriptome (Amrute et al., 2022; Burdziak et al., 2023; Cui, Wang, Maan, & Wang, 2023; Erbe, Stein-O’Brien, & Fertig, 2022; Hyttinen, Eberhardt, & Hoyer, 2012; Jiang et al., 2023; Kamimoto, Stringa, et al., 2023; J. Lambert et al., 2023; Osorio et al., 2022; Qiu et al., 2022; Roohani, Huang, & Leskovec, 2022; Theodoris et al., 2023; Tran, Yang, Yang, & Ormerod, 2022; Wang et al., 2022; Yeo, Saksena, & Gifford, 2021). These expression forecasting methods aim to serve as a new type of general-purpose screening tool. Compared to Perturb-seq and similar assays, *in-silico* modeling is cheaper, less labor-intensive, and easier to apply to less accessible cell types (Bunne et al., 2024). Applications frequently include nomination, ranking, or screening of up to hundreds of genetic perturbations that are expected to have interesting or valuable effects on cell state (Burdziak et al., 2023; Erbe et al., 2022; Jiang et al., 2023; Kamimoto, Stringa, et al., 2023; Roohani et al., 2022; Theodoris et al., 2023; Wang et al., 2022; Yeo et al., 2021). For example, expression forecasting is being used to optimize reprogramming protocols; to search for anti-aging transcription factor (TF) cocktails; and to nominate drug targets for heart disease (Amrute et al., 2022; Kamimoto, Adil, et al., 2023; Kimmel, 2024; Theodoris et al., 2023). In many contexts, expression forecasting is poised to augment or even circumvent genetic screens as a candidate gene selection method.

Existing scholarship provides a number of reasons to believe that expression forecasting under novel genetic perturbations can work, but empirical tests lag behind available methods and data. Reasons for optimism include mathematical identifiability guarantees (Lopez, Hütter, Pritchard, & Regev, 2022; Yeo et al., 2021), prior knowledge of gene function (Roohani et al., 2022), catalogs of TF-to-target binding from ChIP or motif analysis (Burdziak et al., 2023; Kamimoto, Stringa, et al., 2023; Tran et al., 2022; Wang et al., 2022), and empirical tests against simulated (Erbe et al., 2022) or real perturbation outcomes (Cui et al., 2023; Jiang et al., 2023; Kamimoto, Adil, et al., 2023; Kamimoto, Stringa, et al., 2023; Roohani, Huang, & Leskovec, 2024; Theodoris et al., 2023; Wang et al., 2022; Yeo et al., 2021). However, existing empirical results have shortcomings. Genetic perturbation encompasses a vast range of possible experiments: for example, PSC reprogramming via overexpression of TF’s yields a completely different predictive task compared to knockdown of stress-response proteins in K562 cells. Whereas data from diverse contexts will be crucial to build general-purpose models of the cell (Bunne et al., 2024), most expression forecasting methods have been tested in only a small number of cellular contexts (Cui et al., 2023; Jiang et al., 2023; Lopez et al., 2022; Roohani et al., 2022; Theodoris et al., 2023; Tran et al., 2022; Wang et al., 2022). Expression forecasting benchmarks on mammalian data have often been conducted by the authors of competing prediction tools in a process that iterates between running tests and altering or tuning methods.

This is necessary for method development, yet it raises the possibility of over-optimistic results due to unknown experimenter degrees of freedom (Boulesteix, Lauer, & Eugster, 2013; Nießl, Herrmann, Wiedemann, Casalicchio, & Boulesteix, 2022). Because benchmarks are so labor-intensive and compute-intensive, direct comparisons between methods on designated held-out data have been limited (Ahlmann-Eltze, Huber, & Anders, 2024; Wu et al., 2024). Interpreting forecasts and improving tools will require systematic benchmarking of expression forecasting performance on a diverse collection of data.

Here, we fill this gap by creating a systematic expression forecasting benchmarking platform that enables neutral evaluation across varied methods, parameters, and datasets. Achieving this required three distinct components, which we have developed so that they are easily reusable and extendable beyond this study. First, we assembled, quality-controlled, and uniformly formatted 11 perturbation transcriptomics datasets (Adamson et al., 2016; Dixit et al., 2016; Frangieh et al., 2021; Freimer et al., 2022; Joung et al., 2023; Nakatake et al., 2020; Norman et al., 2019; Replogle et al., 2020, 2022). Second, for many expression forecasting methods, a key design element is a sparse, interpretable gene regulatory network (GRN) inferred from a combination of motif analysis and gene expression data. Therefore, we assembled a uniformly formatted collection of cell type-specific gene networks derived from motif analysis, co-expression, and other approaches (Cahan et al., 2014; Gerstein et al., 2012; Kamimoto, Adil, et al., 2023; Lachmann et al., 2010; Marbach et al., 2016; Pierson et al., 2015; Xu et al., 2021). Finally, we created a software suite called PEREGGRN (“PErturbation Response Evaluation via a Grammar of Gene Regulatory Networks”, pronounced “peregrine”). Introducing this grammar allows us to flexibly combine features of different possible GRN inference methods, such as different regression methods, different causal network structures, or different ways of handling time (File S1). Our benchmarking software is also configurable, allowing users to easily choose different numbers of genes, datasets, data splitting schemes, or performance metrics. Taken together, our software and online documentation allow users to easily specify experiments across a broad range of expression forecasting models and evaluation schemes.

## Results

### Large-scale perturbation datasets display a high rate of success in targeting individual genes and mostly small global effects

As a first step in building a comprehensive benchmarking infrastructure, we collected datasets with transcriptome-wide profiles of many genetic perturbations (Table 1). For clarity, we will always refer to each dataset using the identifier shown in Table 1. We focused on datasets with many genes perturbed and on datasets that were previously used to showcase expression forecasting methods. We focused on human data: despite the abundance of perturbation transcriptomic data in S*. cerevisiae*, *E. coli*, and *C. elegans*, (Alam et al., 2016; Hackett et al., 2020; Han, Li, Filko, Li, & Zhang, 2023; Kemmeren et al., 2014; MacNeil et al., 2015), such data are less useful for drug target discovery or optimizing directed differentiation protocols for stem cells.

**Table 1.** Perturbation transcriptomics data used in this study. OE: overexpression; RNP: ribonucleoprotein; PSC: pluripotent stem cell. The CRISPRa system used by Norman et al. is a dCas9-SunTag fusion that recruits multiple copies of an scFV-VP64 fusion, where scFV is a single-chain antibody and VP64 is a transcriptional activator (Gilbert et al., 2014).

| Identifier | Transcript-omic method | Citation | Genes perturbed | Gene selection | Perturbation method | Dura-tion, days | Cell line |
| --- | --- | --- | --- | --- | --- | --- | --- |
| nakatake | SureSelect, SurePrint G3 (Agilent) | (Nakatake et al., 2020) | 714 | Curated TF's | OE: Transfection with dox-inducible transgenes | 2 | SEES3 (PSC) |
| nakatake scrna simulated: resampled with more samples and less depth per sample (methods). |  |  |  |  |  |  |  |
| joung | 10X 3' scRNA | (Joung et al., 2023) | 3,548 isoforms | All TFs | OE: Lentiviral ORF transduction | 7 | H1 (PSC) |
| norman | 10X 3' scRNA | (Norman et al., 2019) | 112 | Screen for fitness | CRISPRa: see legend | 7 | K562 |
| replogle1 | 10X 3' scRNA | (Replogle et al., 2020) | 63 | Manual | CRISPRa: dCas9-SunTag/scFV-VP64 | 8 | K562 |
| replogle3 | 10X 3' scRNA | (Replogle et al., 2022) | 2285 | All essential genes | CRISPRi: lentiviral delivery of gRNA and dCas9-KRAB | 6 | K562 |
| replogle4 | 10X 3' scRNA | (Replogle et al., 2022) | 2285 | All essential genes | CRISPRi: lentiviral delivery of gRNA, dCas9-Zim3-KRAB | 7 | RPE1 |
| adamson | 10X 3' scRNA | (Adamson et al., 2016) | 87 | UPR-related genes | CRISPRi: dCas9-KRAB and lentiviral gRNA transduction | 7 | K562 |
| replogle2 | 10X 3' scRNA | (Replogle et al., 2022) | 9867 | All expressed genes | CRISPRi: lentiviral delivery of gRNA and dCas9-KRAB | 8 | K562 |
| replogle2 large effect: as above, but controls and perturbations with energy score above 0.6. |  |  |  |  |  |  |  |
| replogle2 tf only: as above, but controls and perturbations of TF's. |  |  |  |  |  |  |  |
| freimer | 3' Tag-seq | (Freimer et al., 2022) | 24 | Regulators of IL2RA, IL-2,CTLA4 | CRISPR: Cas9 RNP with lentiviral gRNA delivery | 5 | CD4+ T cells |
| dixit | 10X 3' scRNA | (Dixit et al., 2016) | 24 | Prior work (Amit et al., 2009) | CRISPR: Cas9 mouse; lentiviral gRNA delivery | 7 | K562 |
| frangieh | 10X 3' scRNA | (Frangieh et al., 2021) | 248 | Checkpoint inhibitor resistance genes | CRISPR: Cas9 RNP nucleofection and lentiviral gRNA delivery | 14 | Primary melano-ma |
| frangieh IFNg v1: IFN-gamma exposed subset, preprocessed identically to DCD-FG experiments |  |  |  |  |  |  |  |
| frangieh IFNg v2: As v1, but with stricter inclusion criteria (methods) |  |  |  |  |  |  |  |
| frangieh IFNg v3: As v2, but pseudo-bulk aggregated within each gRNA combination (methods) |  |  |  |  |  |  |  |

To detect and minimize technical issues, we conducted typical exploratory analysis, quality control, and dataset-specific normalization (Methods). To understand the success rate of the perturbations and the size of the downstream effects, we examined targeted-locus and transcriptome-wide effect size (Fig 1A,B). As a measure of robustness, we examined the Spearman correlation in log fold change between replicates. In cases lacking biological replicates, we examined correlation between technical replicates (e.g. different guide RNA’s) or the number of differentially expressed genes. The datasets contain a minority of perturbations with very large effects, and the larger effects tend to have higher correlation between replicates. For each overexpression or knockdown experiment, the targeted genes’ expression mostly increases or decreases as expected, with the lowest rate being joung, nakatake, and replogle1: in joung, 73% of overexpressed transcripts increased as expected, and in nakatake and replogle1, about 92% of overexpressed transcripts increased as expected. In knockout experiments, cells with a nonsense mutation or a truncated sequence could in principle express high levels of detectable mRNA while experiencing complete loss of function of the encoded protein, but we still observe that target gene RNA levels frequently decrease. In knockdown and overexpression experiments, we removed samples where the targeted transcript does not decrease or increase as expected. In knockout experiments, we retained all samples except those flagged for quality issues by the original authors. For adamson, dixit, and norman, we eschew these filters and instead use preprocessing identical to the GEARS evaluations (Roohani et al., 2024). For frangieh, we again eschew filtering on targeted-gene effect. We preprocess frangieh data in multiple ways, and version 1 is identical to that used by the DCD-FG evaluations (Lopez et al., 2022). A full accounting of our preprocessing is given in Methods. In summary, this collection provides conservative quality control and a well-characterized testing ground for expression forecasting under genetic perturbation.

**Fig 1.**
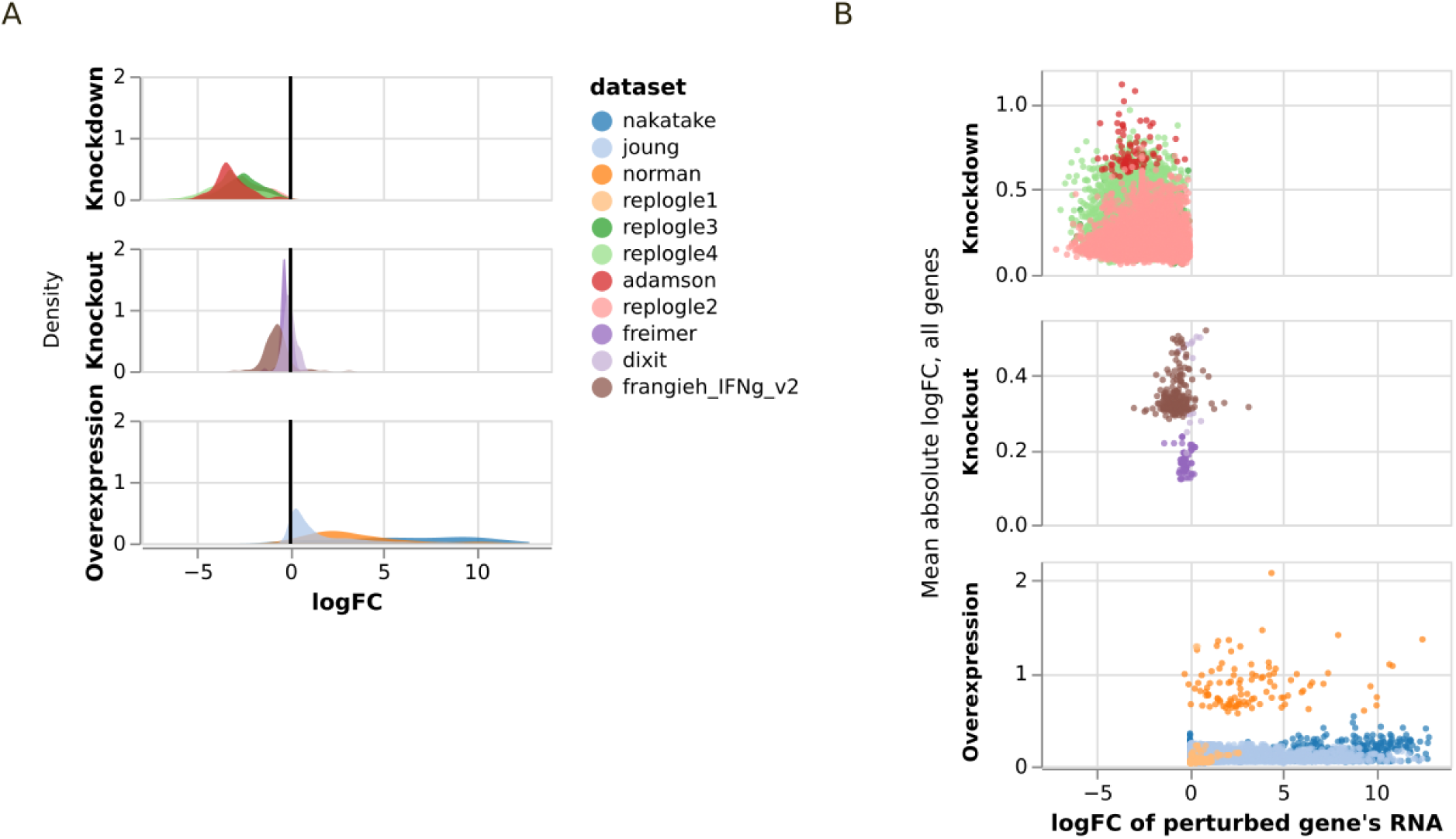
Quality analysis of large perturbation datasets. All values shown in this figure are computed after filtering, aggregation, and normalization as described in the Methods. A. Effect on total-count-normalized transcript levels of the perturbed gene (X-axis). B. Effect on the perturbed gene (X-axis) and the whole transcriptome (Y-axis). Log fold change (logFC) over controls is estimated using a pseudocount of 1. For norman, dixit, and adamson, only 6,000 highly variable genes are included.

In most of these datasets, genes to perturb were not selected at random, but rather were known to be important for survival or regulation (Table 1). One exception is replogle2, in which all expressed genes were knocked down. To understand how non-random perturbation selection affects results, we separately labeled subsets of replogle2 that include 1) only TF perturbations or 2) only large effects.

### Regression-based analyses do not out-perform non-informative baselines

Our first set of experiments are inspired by the modular structure of CellOracle (Kamimoto, Stringa, et al., 2023). Starting with a causal structure derived from motif analysis, CellOracle trains ridge regression models to predict each gene from its candidate regulators (Kamimoto, Stringa, et al., 2023). Many expression forecasting methods are based on different network structures or different supervised machine learning models (Burdziak et al., 2023; Erbe et al., 2022; Kamimoto, Stringa, et al., 2023; Lopez et al., 2022; Qiu et al., 2022; Wang et al., 2022). It is natural to ask what regression methods and network structures perform best, while holding other factors constant.

Existing software is typically oriented towards end-to-end solutions, with limited flexibility to replace individual pipeline components. CellOracle allows user-defined network structures, but it offers only two alternative regression methods, and it is not designed to be trained on interventional data. To train models on interventional data using different network structures and regression methods, we created a Grammar of Gene Regulatory Networks (GGRN). GGRN flexibly combines features of different possible GRN inference methods, such as different regression methods, different causal network structures, or different ways of handling time (File S1). The specified regression method is trained to predict the log-scale expression of each target gene using the log-scale expression of all TF’s as features (excluding the target gene if it is a TF). Importantly, samples where gene *j* is directly perturbed are omitted when training models to predict gene *j*’s expression.

To predict perturbation outcomes, we begin with the average expression of all controls. The perturbed gene is then set to 0 (for knockout experiments) or to its observed value after intervention (for knockdown or overexpression experiments). The resulting expression values are provided to the regression models as features, and the corresponding regression model predictions are what we evaluate. A key aspect of our evaluations is a non-standard data split: no perturbation condition is allowed to occur in both the training and the test set. Randomly chosen perturbations and all controls are allocated to the training data, while a distinct set of perturbations is allocated to the test data. Performance on held-out perturbations is essential to understand these tools’ potential for *in silico* perturbation screening (Fig. 2).

**Figure 2.**
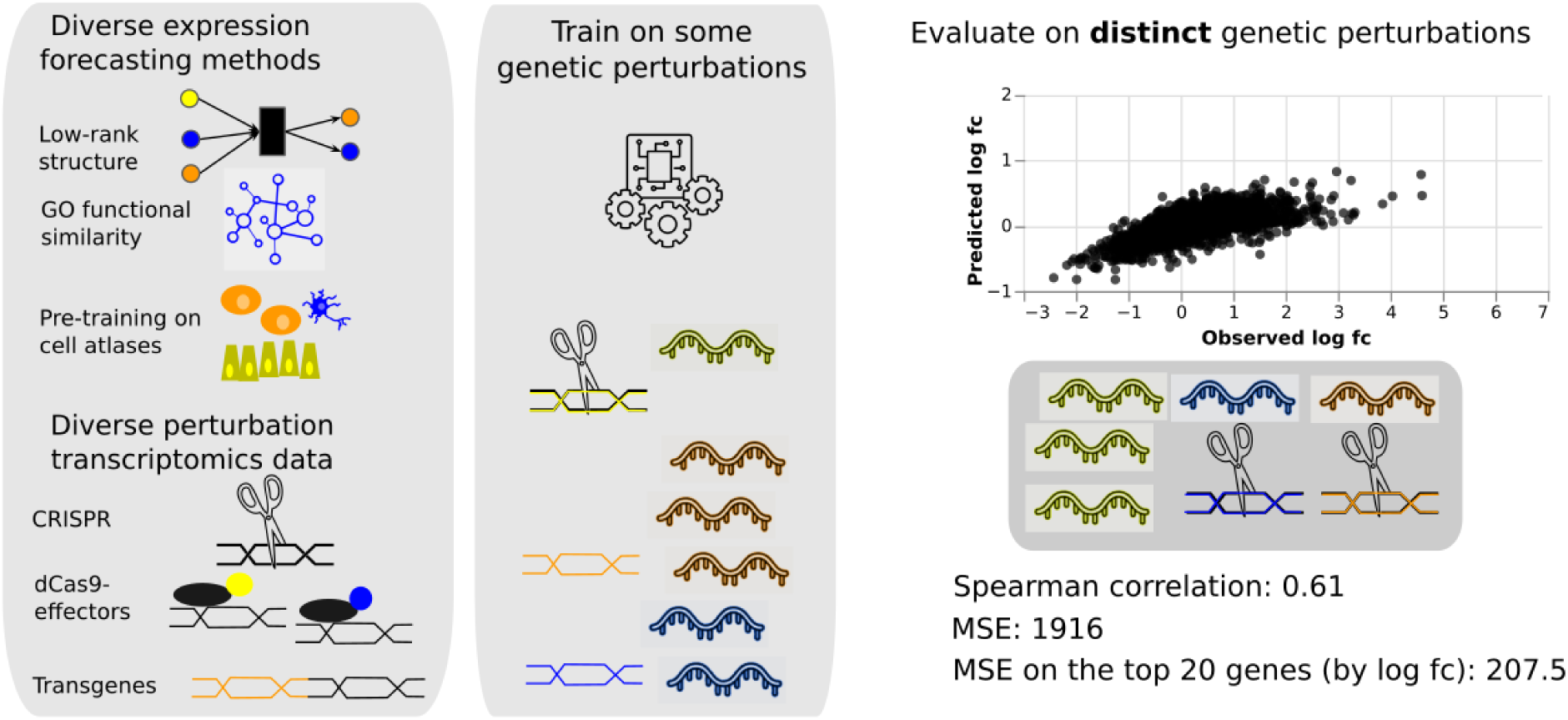
Evaluation plan. We created a uniform interface to a diverse collection of methods for GRN inference. We collected perturbation transcriptomic data from diverse human cell types and genetic perturbation methods. Data are partitioned into a training set and a test set such that no perturbation is present in both sets. Predicted and observed log fc over controls are compared via several performance metrics to determine which biological systems, types of input data, computational methods, or characteristics of genes are associated with successful prediction of expression fold change under the novel perturbations. This figure uses clipart from the Noun Project: “crispr” by Cornelia Scheitz and “rna” by Bacontaco, each licensed under [CCBY-SA 4.0](https://creativecommons.org/licenses/by-sa/4.0/), and “Machine Learning” by apixlabs and “Network” by Meaghan Hendricks, each licensed under CC BY-SA 3.0.

We computed a variety of evaluation metrics (Table 2). These methods fall into three broad categories. Some are obvious or commonly used performance metrics (mean absolute error (mae), mean squared error (mse), spearman correlation, proportion correct direction). Others are computed on the top 100 most differentially expressed genes, emphasizing signal over noise for datasets with sparse effects. Finally, accuracy when classifying cell type is of special interest in studies of reprogramming or cell fate. Notably, some methods are intended to provide only cell-type classifications or low-dimensional embeddings (Cui et al., 2023; Kamimoto, Stringa, et al., 2023; Theodoris et al., 2023; Yeo et al., 2021), while others provide gene-specific predictions of expression, velocity, or fold-change (Burdziak et al., 2023; Erbe et al., 2022; Qiu et al., 2022; Roohani et al., 2022; Wang et al., 2022). We will show that different metrics may give substantially different conclusions.

**Table 2.**
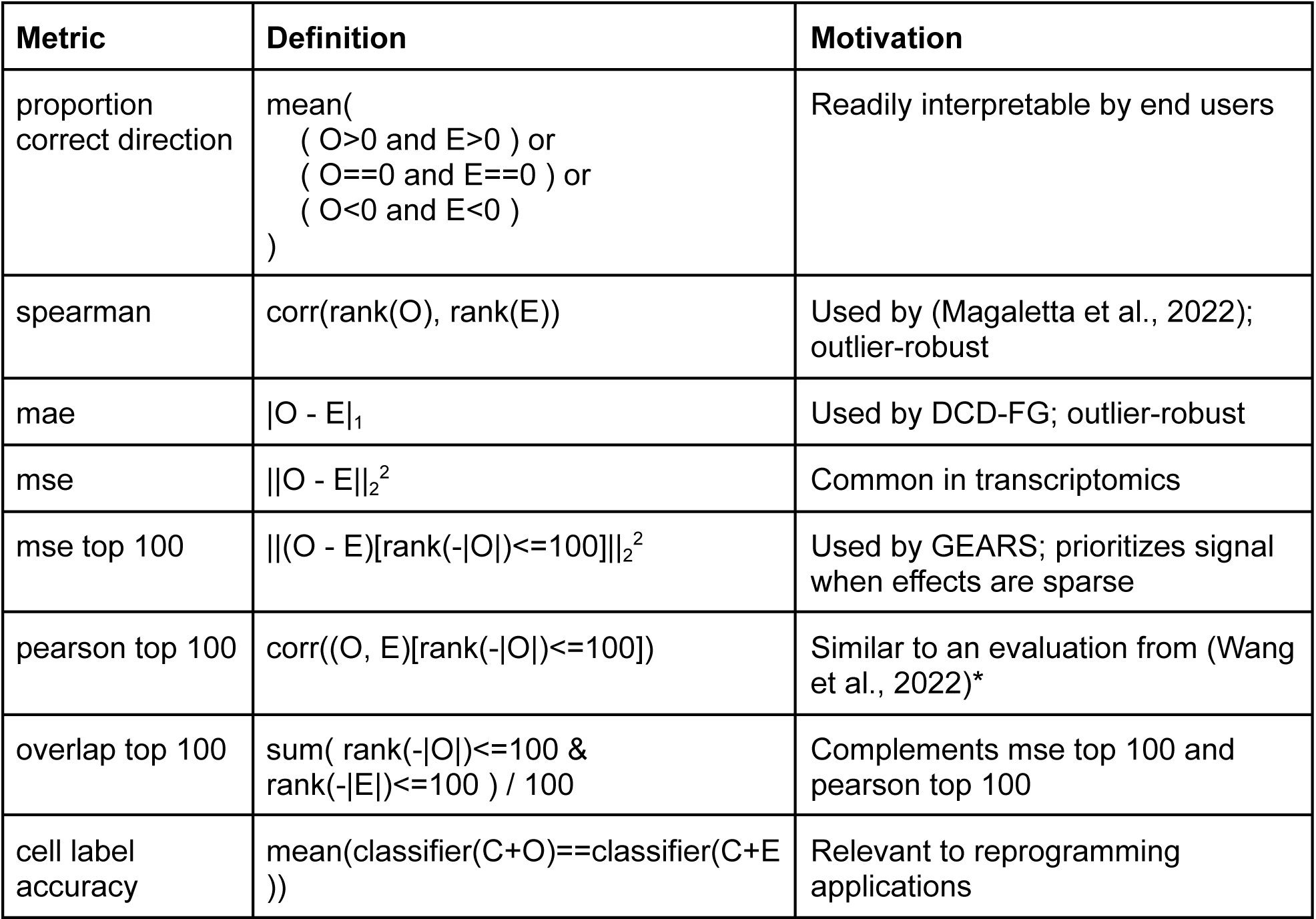
Evaluation metrics. O indicates observed log fold change, E indicates expected log fold change, and C indicates log-scale expression of controls. Classifiers are trained on Louvain cluster labels in the training data (Methods). *Wang et al. use a p-value cutoff, meaning their evaluation can include a variable number of genes depending on the observed differential expression, not always the top 100 genes.

We compared nine supervised learning methods. In contrast to EF evaluations to date, we also included mean and median baseline predictors in which ignore the input features and return the training-set mean or median of the target (dependent) variable. We further included a “linear embedding” baseline method based on linear functions of gene embeddings that showed good relative performance in a recent benchmark study (Ahlmann-Eltze et al., 2024). To our surprise, the mean or median baseline was always the top performer (Fig. 3, left column). Some performance improvements over mean and median baselines were seen on nakatake and replogle3 when evaluating cell label accuracy (Fig. S1, left column), but for the large majority of evaluation metrics and datasets, mean and median baselines performed best.

**Figure 3.**
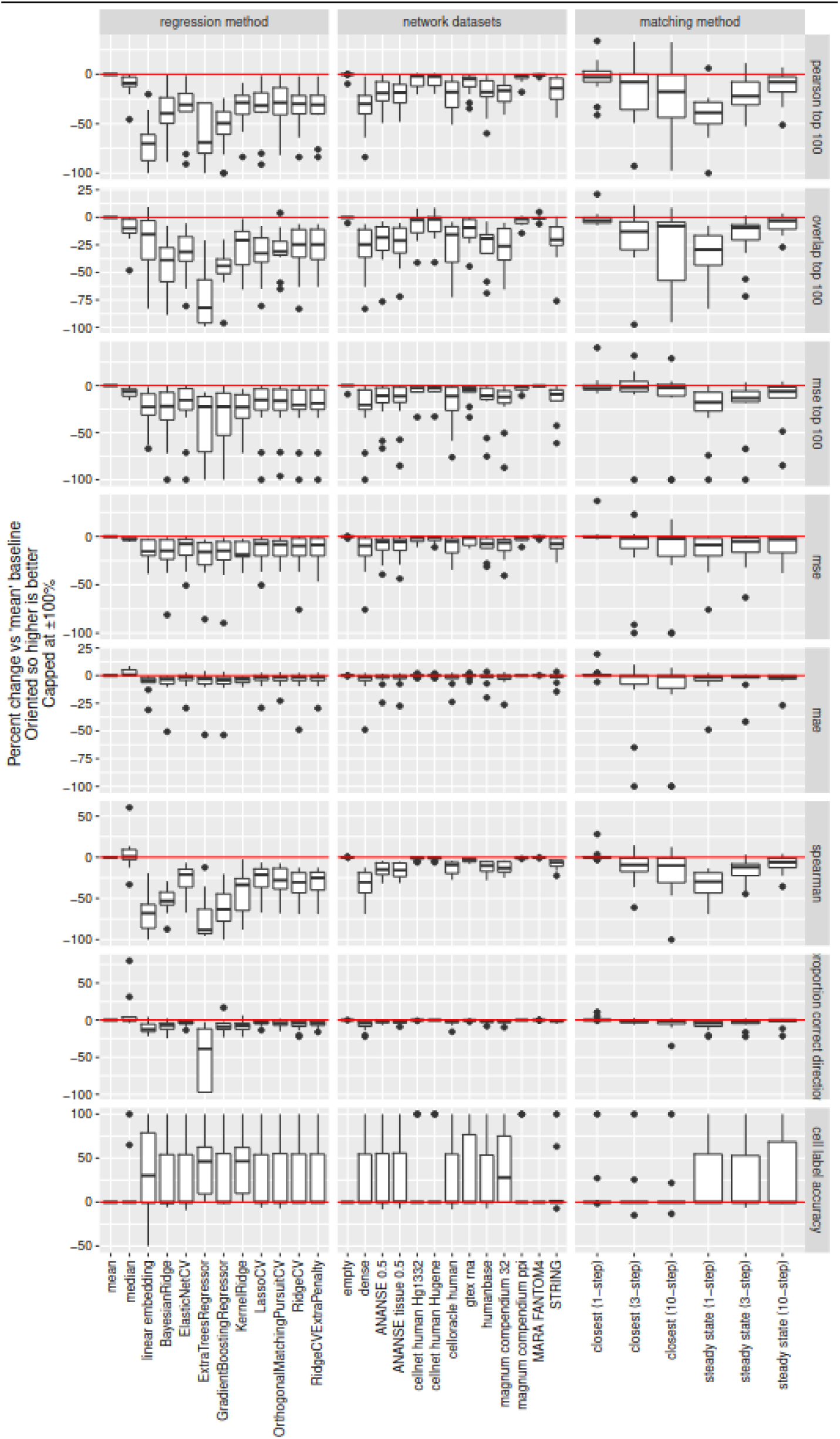
Evaluation of supervised ML and feature selection methods against simple baseline predictors. Each dot represents performance of one method on one dataset by one metric. Boxplots summarize performance across all datasets. The right margin shows the performance metrics described in Table 2. The y axis indicates performance as the percent change over the ‘mean’ baseline, capped at ±100%. Metrics where larger values would normally indicate worse performance, such as mse, are inverted such that percentages above zero always indicate improved performance. The x axis labels describe which regression method was used (left), which network was used for feature selection (middle), or how models handle time (right). “Closest” indicates that features used to predict each perturbed profile are drawn from the closest control profile, except for the perturbed gene itself. “Steady state” indicates that features used to predict each perturbed profile are drawn from that same profile.

**Figure S1.**
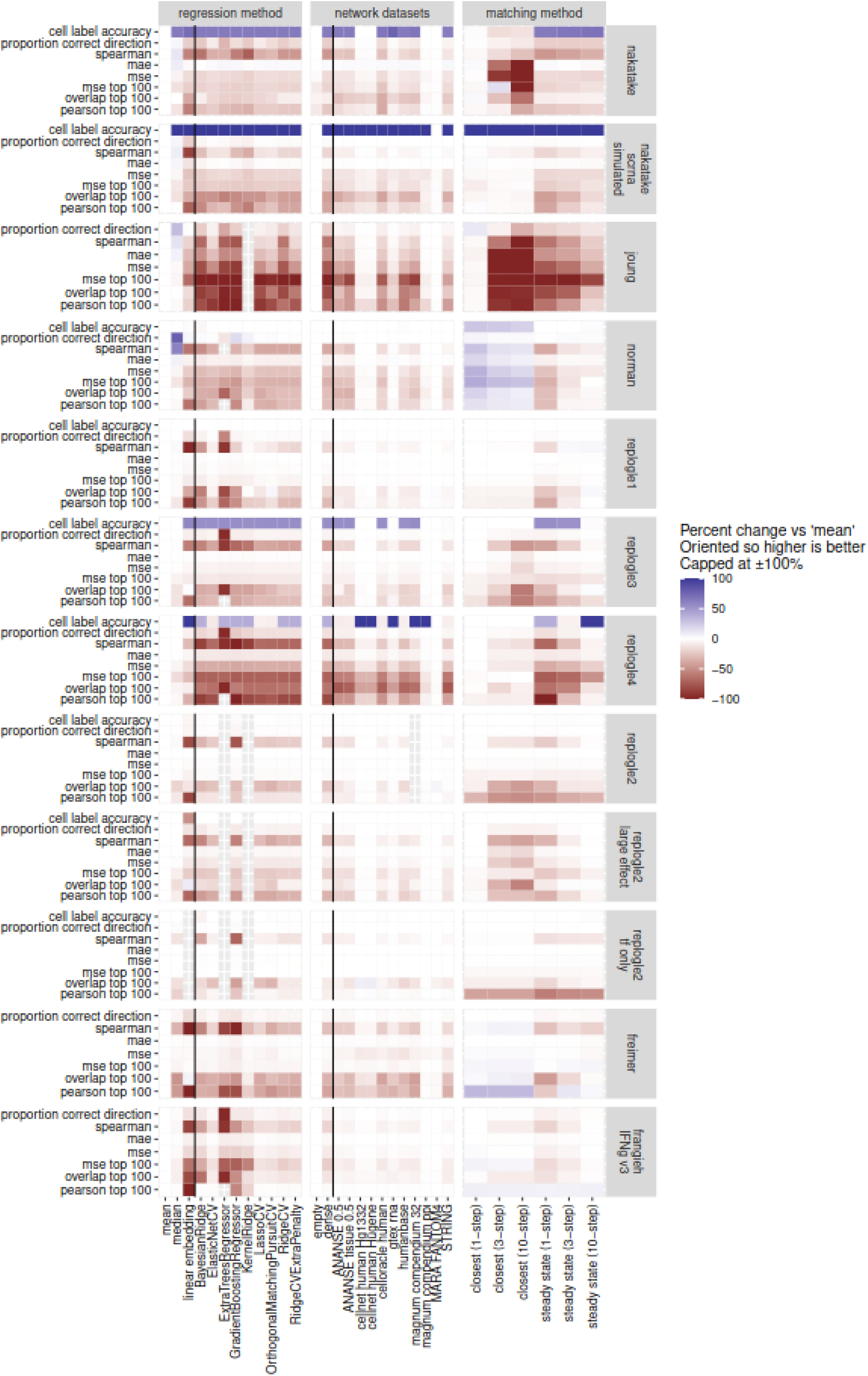
Related to Fig. 3. Various performance metrics to assess prediction of held-out perturbations. All experiments within each dataset use the same data split and can be directly compared. The y axis shows different performance metrics as described in Table 2. The x axis labels describe which regression method was used (left), which network was used for feature selection (middle), or how models handle time (right). Black lines separate baselines within each panel. In the right column, “closest” indicates that features used to predict each perturbed profile are drawn from the closest control profile, except for the perturbed gene itself. “Steady state” indicates that features used to predict each perturbed profile are drawn from that same profile. ExtraTrees and KernelRidge are missing from certain panels due to memory requirements exceeding 64GB. The Ahlmann-Eltze baseline is missing from the replogle2 tf only panel because the entire training set consisted of genes that were perturbed but not measured in the original source data.

We suspected these generally negative results could be driven by overfitting or lack of causal identification, so we sought to reduce the difficulty of the task. In several common cell lines, TF regulons based on evidence of direct binding are enriched for target genes that respond to perturbation of those TFs (Minaeva, Domingo, Rentzsch, & Lappalainen, 2024), and many methods restrict causal relationships by predicting a target gene’s expression using only TF’s with nearby motif occurrences in promoters or enhancers (Kamimoto, Stringa, et al., 2023; Pemberton-Ross, Pachkov, & van Nimwegen, 2015; Wang et al., 2022; Young, Raftery, & Yeung, 2014). This yields easier regression problems with far fewer predictors. We collected 10 sets of previously published gene networks that had been generated in a variety of ways, including co-expression analysis, shared function of gene products, and cis-regulatory element motif analysis (Table 3). We processed them into a uniform format and incorporated them into our benchmarking software (Methods). The software allows users to easily train diverse supervised ML methods using expression of network neighbors as the input features.

**Figure S2.**
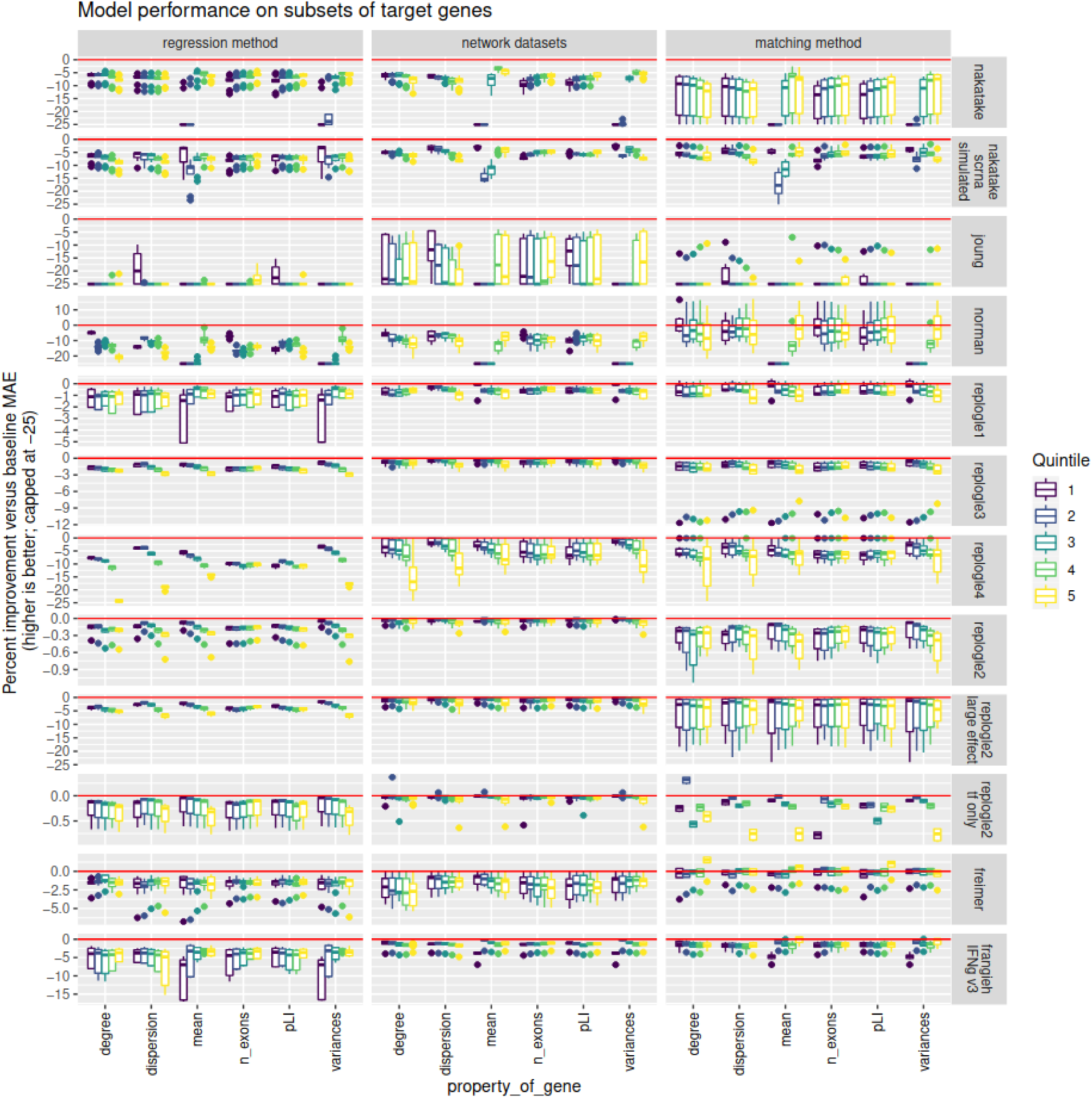
Related to Fig. 3. Grouping target genes into quintiles using gene-level features reveals worse-than-baseline mae for most gene sets. The y axis shows the mae contribution of each gene set, expressed as percent change from the best baseline (inverted so that higher is better). The x axis indicates which feature we stratified genes on, and within each feature, the color indicates the quintile. The figure includes 6 of the 45 features included in this analysis.

**Table 3.**
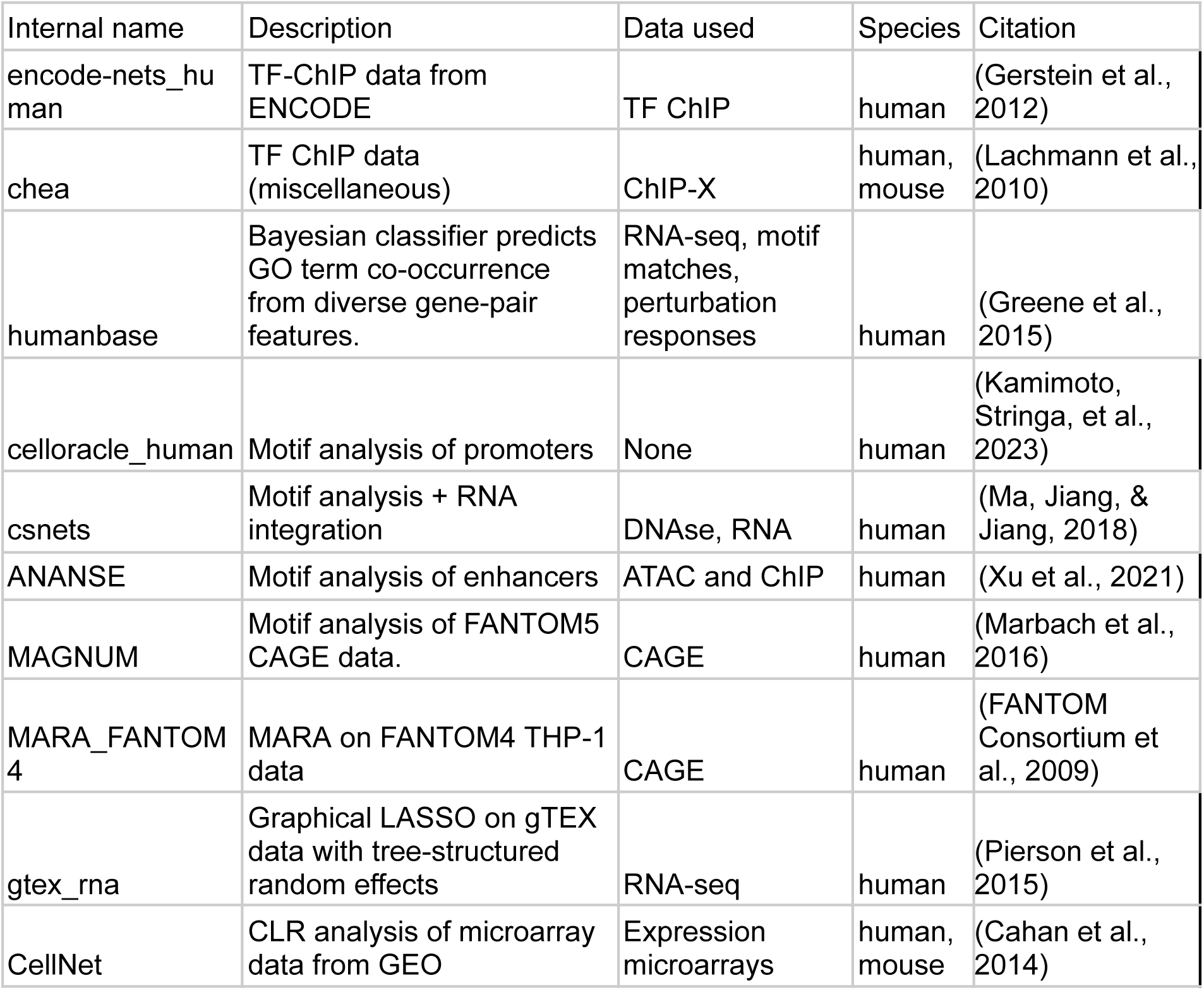
Published networks that are expected to be enriched for human transcriptional regulatory relationships.

Separately using each of several networks for feature selection, we trained ridge regression models with TF expression as features and target gene expression as labels. (Ridge regression is a simple, fast approach with previous success in interpretable models for post-perturbation expression forecasting (Kamimoto, Stringa, et al., 2023).) We predicted expression after held-out interventions, then evaluated performance. As controls, we included a dense network (all features are used for each target) and an empty network (no features are used, equivalent to the “mean” baseline). These results use the same data split as the regression model tests described above, so performance can be directly compared. Use of certain prior network structures led to improved cell type accuracy, but by other metrics, the mean, median, or empty network always remained among the top performers (Fig. 3 and Fig. S1 center column).

Finally, we conducted additional experiments surrounding handling of time, another feature of regression-based methods whose treatment in the literature varies. The simplest way of training a regression model for expression forecasting uses predictive features drawn from the same cell as the target (dependent) variable. Viewing the transcriptome as a dynamic process, this implies an assumption that the regulation of interest can be inferred from steady-state measurements alone (Kamimoto, Stringa, et al., 2023; Lopez et al., 2022; Wang et al., 2022). But some tools do not assume steady state, rather predicting gene expression from features drawn from a control sample or a previous time point, with only the perturbed gene(s) altered (Burdziak et al., 2023; Roohani et al., 2022; Yeo et al., 2021). Among steady-state methods, some tools iterate predictions to simulate total effects, and the number of iterations can affect the content and quality of the results (Kamimoto, Stringa, et al., 2023; Wang et al., 2022). To understand the effect of these choices, we constructed features to predict each observation from either itself (“steady-state” in Fig. 3 and Fig. S1) or from the closest control (“closest” in Fig. 3 and Fig. S1). When controls were chosen, we set the perturbed gene to its post-perturbation expression level. We iterated predictions for one, three, or ten time-steps. These configurations did not outperform baselines overall (Fig. 3), but did succeed on certain datasets and metrics (Fig. S1, right column).

### Very few subsets of genes are predictable

Even if expression forecasting methods do not outperform baseline predictors in terms of transcriptome-wide mean absolute error, they may work for certain target genes or perturbations. The mae has the advantage of being additive, so contributions from individual samples or features can be examined separately. We used this additivity to search for predictable sets of genes, which we define as sets of genes whose contribution to mae is lowest for a non-baseline model. We grouped target genes into quintiles of 45 gene-level features, including probability of being loss-of-function intolerant (Lek et al., 2016); number of exons; mean, variance, or dispersion in the training data; and in-degree or out-degree from 10 published networks. This yielded 64,563 total comparisons across 12 datasets, 26 non-baseline inference methods, 45 factors to stratify genes by, and 5 quintiles per factor, with some combinations absent (Fig. S2). Of these gene sets, 2,071 were predictable (an informative model outperformed all baselines), and 62,492 were not predictable (a baseline model was best). Above-baseline performance was concentrated on matching-method experiments on the Norman data (Fig. S2). When grouping by perturbed gene instead of target gene, zero out of 62,928 comparisons showed an informative model outperforming all baselines. Thus, poor performance of simple expression forecasting methods is not driven by specific categories of genes, and it is also not avoidable by focusing on any specific subset of genes.

### Investigating strong performance of the “mean” baseline

We next sought to understand why the mean of the training data has such strong relative performance. Indeed, the “mean” baseline works surprisingly well even in absolute terms: the Spearman correlations between the observed and predicted log fold changes are typically well above zero (Fig. S3a). Recall that because the fold change is defined relative to control experiments, the mean is not expected to be zero, and does vary from gene to gene. To explain the performance of the mean baseline, we considered two hypotheses that are not mutually exclusive: a statistical hypothesis rooted in the bias-variance tradeoff and a biological hypothesis rooted in stereotypical responses of each gene to perturbation. Here, we demonstrate that either could contribute to the surprisingly strong mean baseline performance we observe.

Regarding the bias-variance tradeoff, consider two alternative estimators of baseline expression: the mean of all training data or the mean of only the control samples. The mean of the control samples is unbiased, but given limited sample sizes, it has high variance. The mean of all training data will be biased by perturbation effects, but will have lower variance. If perturbation effects are generally small and noise levels generally high, the mean of all samples may be a better estimator of baseline expression. A small simulation (Methods) shows that in simulated data with only noise and no true perturbation effects, the mean of the training data can easily yield correlations greater than 0.6 between observed and predicted expression changes. This simulation illustrates how the “mean” baseline can yield large, positive correlations between predicted and observed log fold change over controls.

Alternatively, we consider a second hypothesis: stereotypical perturbation responses. A stereotypical response for a given gene is similar across many perturbations, perhaps due to common affected signaling pathways, cell stress, or inadequate controls (e.g. off-target effects of a control guide RNA). If perturbation responses are stereotypical, the mean of the training data would perform well on held-out perturbations as it learns the usual response pattern of each gene. Of course, there may be perturbations for which a gene deviates from its stereotypical response, but there is still information in the mean. Specifically, this hypothesis predicts that for independent experiments A and B, fold change of treatment A over control A will be correlated with fold change of treatment B over control B, even when control A and control B are generated independently and even when treatment A is different from treatment B. For datasets with replicated controls, we estimated this type of correlation for 100 independent pairs of treatment-over-control log fold change. It was positive and far from zero for some datasets, notably joung, replogle4, replogle2 large effect, and nakatake (Fig. S3B). The same datasets tended to have the highest correlations between predicted and observed log fold change in the “mean” baseline analyses (Fig. S3A).

**Figure S3.**
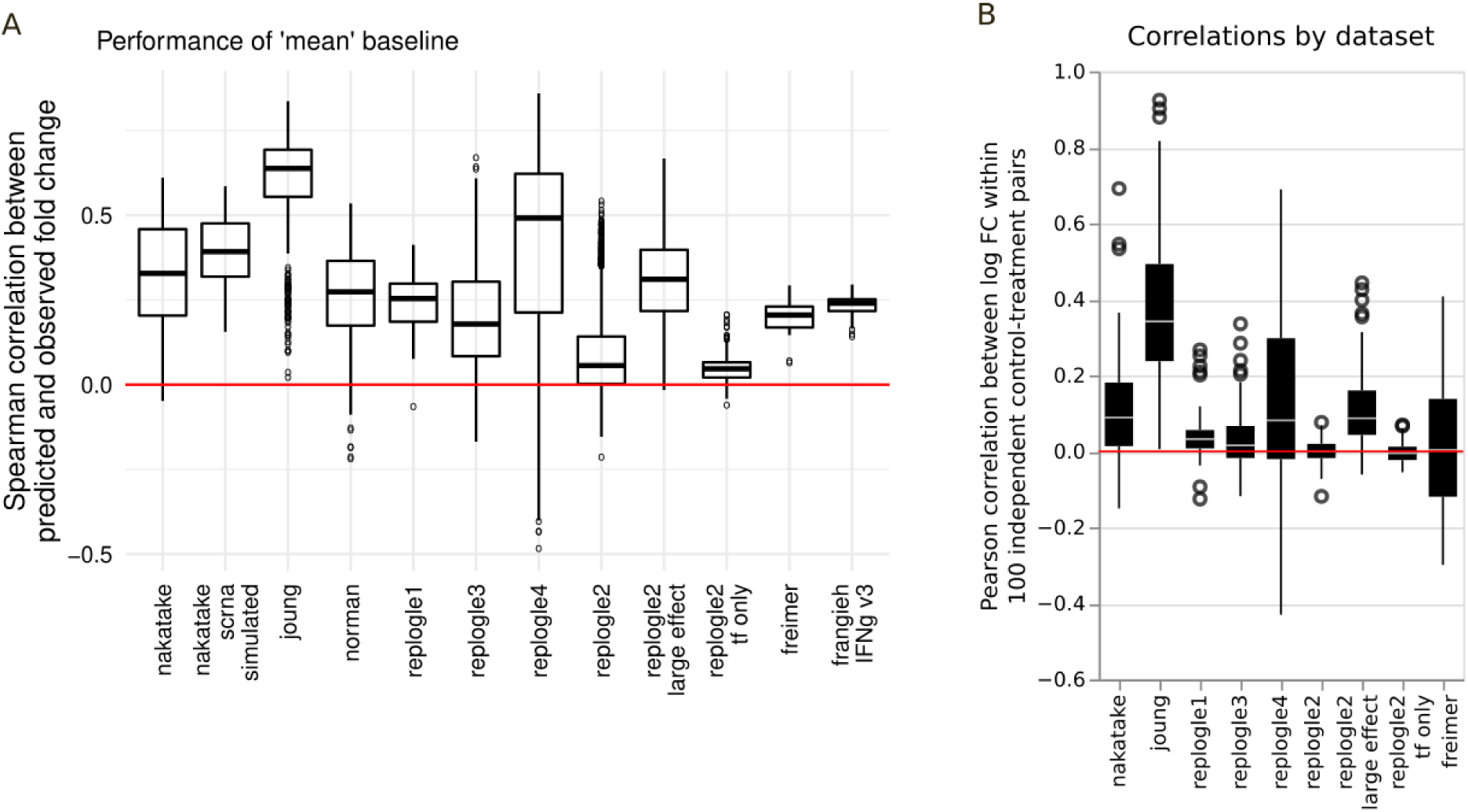
Investigating strong performance of the “mean” baseline. A. Spearman correlation between observed fold change over controls and fold change over controls predicted by the “mean” baseline (y axis) across various datasets (x axis). B. Pearson correlation between observed fold change of treatment A over control A and treatment B over control B, where treatments A and B are different perturbations and controls A and B are separate replicates. Display includes 100 randomly selected pairs per dataset.

### In network-based simulated data, correct network structures perform best

In some mathematical formulations, GRN reconstruction requires all genes to be perturbed, and held-out perturbations cannot necessarily be predicted (Hyttinen et al., 2012; Wagner, 2001; Zhang et al., 2023). To determine whether evaluation on held-out perturbations can yield meaningful results in an ideal scenario, we simulated data from autoregressive models whose structures were based on several known networks (Methods). Compared to a fully connected network, an empty network, or another mis-specified network, the network used to generate the data achieves equal or slightly better performance (Fig. S4). Other top performers were often networks from similar sources (Fig. S4). This shows that correctly specified structural models can successfully predict responses to unseen perturbations.

**Figure S4.**
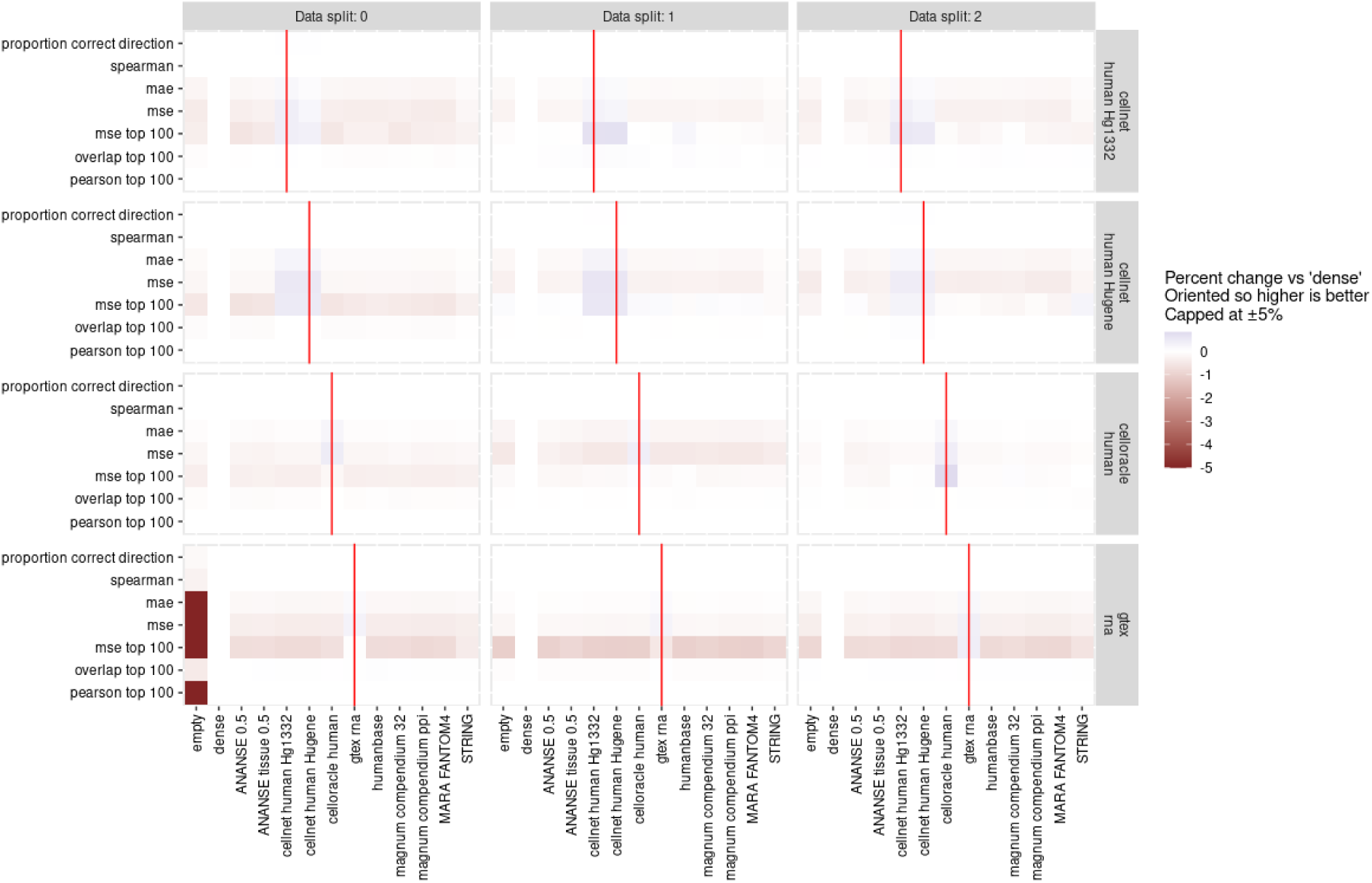
Related to Fig. 3. Impact of regulatory network source (x axis) on EF performance (color scale). Evaluations use multiple train-test splits (top margin) of data generated from a specific network structure (right margin). Metrics are defined in Table 2. Data are generated from autoregressive models with random initial state that are run for a single time-step, with matched controls for each sample provided during training (Methods). Red lines indicate that the network used for inference matches the network used to generate the data.

### Review and benchmarking of diverse published methods

Given the preceding largely negative findings, we sought to test additional, previously published expression forecasting methods. We included any computational method capable of predicting the outcome of novel genetic perturbations as long as the training requires only single-timepoint transcriptome data (not time-series data or RNA velocity data). We included all methods with peer-reviewed paper and code available as of Sept 1, 2023, which were DCD-FG, GEARS, and GeneFormer (Lopez et al., 2022; Roohani et al., 2022; Theodoris et al., 2023).

Although all of these methods demonstrate perturbation predictions after training on expression data, they employ very different strategies and they have different primary purposes. DCD-FG is intended for causal graph structure inference, not prediction. GEARS is built for predicting post-perturbation gene expression, but with a primary emphasis on novel genetic interactions rather than unseen interventions. GeneFormer is a general-purpose foundation model with usage demonstrations focusing on gene dosage sensitivity, chromatin dynamics predictions, gene network dynamics predictions, and prediction of cell state after genetic perturbation. Our evaluations are modeled after a specific demonstration in which GeneFormer predicted changes in cardiac muscle expression state in response to genetic perturbation, after being fine-tuned to predict cell type labels but with no fine-tuning on any perturbation data (Theodoris et al., 2023).

When we computed the median performance relative to a non-informative baseline, no methods were consistently better by any metric (Fig. 4). Seeking to reconcile this negative overall result with prior work, we sought to understand how each dataset, evaluation metric, and other specific choices affects the outcomes (Fig. S5, S6).

**Figure 4.**
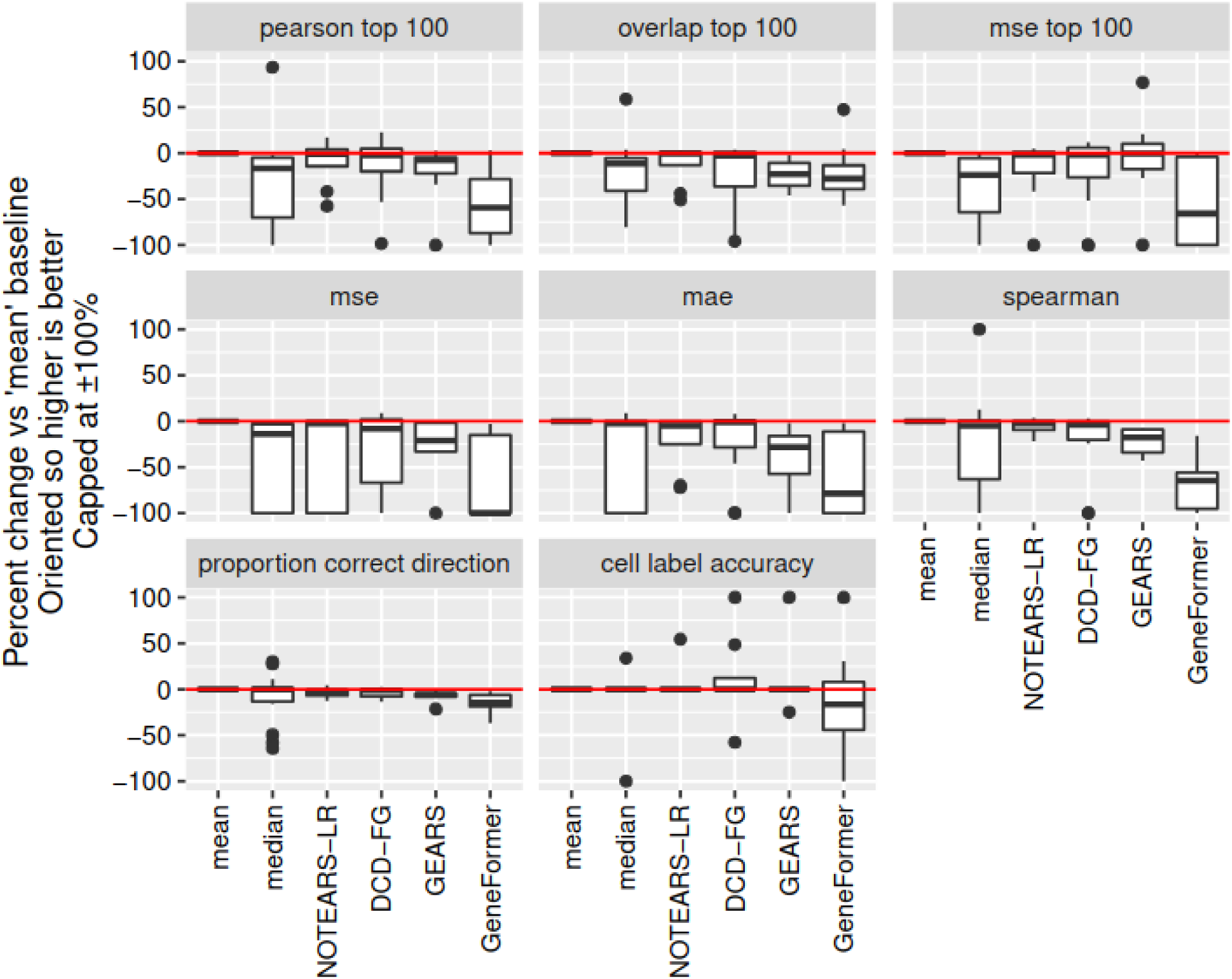
Median performance of published expression forecasting methods. The y axis shows different performance metrics as described in Table 2. The x axis labels describe which method was used. The y axis indicates performance as the percent change over the “mean” baseline. Each boxplot aggregates over all datasets, and each outlying point indicates a single dataset. Variables where larger values indicate worse performance are inverted such that percentages above zero always indicate improved performance.

Using code published by the DCD-FG authors, we repeated the benchmarks of DCD-FG on the gamma interferon-treated subset of frangieh, which was used for expression forecasting benchmarks in the DCD-FG paper (Lopez et al., 2022). Comparisons are made against NO-TEARS-LR and NO-TEARS, which are closely related to DCD-FG and are also implemented in the DCD-FG software (Zheng, Aragam, Ravikumar, & Xing, 2018). The relative performance of DCD-FG and related methods is consistent with Lopez et al. (Fig. S5a). In addition to the methods implemented in DCD-FG, we included an IID Gaussian baseline that ignores perturbations, and we fit either a diagonal or a full covariance matrix. Surprisingly, the IID baseline outperformed all causal inference methods when using a full covariance matrix (Fig. S5a).

When predicting a target gene’s expression, the DCD-FG benchmarks allow each method to use held-out expression of genes that are causally upstream according to the learned model (Romain Lopez, personal communication). This is a natural choice due to the mathematical formulation of DCD-FG, but it does not fit the use-case motivating our study, in which held-out perturbations are completely unobserved. Thus, we modified our benchmarking framework to include DCD-FG and to explicitly control the use of held-out gene expression during prediction. Using the top (roughly) 1,000 genes in each dataset with a gene-selection procedure identical to the DCD-FG paper (Methods), we compared DCD-FG against NO-TEARS-LR and two baselines by a variety of metrics. Different metrics yielded substantially different results in many cases (Fig. S5b). By some metrics, especially mse top 100, DCD-FG and/or NO-TEARS-LR very narrowly outperformed baselines on nakatake, norman, replogle4, and replogle2 large effect (Fig. S5b). However, the largest of DCD-FG’s improvements over baseline mostly required access to held-out expression of upstream genes (Fig. S4b).

Since the DCD-FG model does not include measurement error and the melanoma data contain many perturbational regimes with only one cell measured (66% of regimes), measurement error may substantially affect results. We repeated the frangieh benchmarks after removing perturbational regimes with fewer than 50 cells (frangieh IFNg v2). This preprocessing also eliminated combinations of knockouts and focused the evaluation entirely on perturbations of genes that are not perturbed in the training data, rather than novel combinations of genes that may be perturbed individually in the training data. On this dataset, DCD-FG did not out-perform baselines, even with access to held-out expression of upstream genes (Fig. S5b). To further understand the effect of bulk versus single-cell measurement error, we repeated the same experiments after removing perturbational regimes with fewer than 50 cells and averaging expression within each perturbational regime (this is labeled frangieh IFNg v3). DCD-FG did not out-perform baselines, even with access to held-out expression of upstream genes.

Since Lopez et al. tested DCD-FG on single-cell data, not bulk data, we also repeated the nakatake experiments after simulating single-cell data based on the real bulk transcriptome data (Methods). We further tested DCD-FG on norman, where the focus is on genetic interactions (Norman et al., 2019). In norman, there is a mix of individual perturbations and combinations, so even though no combination is present in both the training and test data, the same gene may be perturbed in both the training and the test data. In both of these experiments, and without having access to post-perturbation expression of upstream genes, DCD-FG’s predictions narrowly outperformed NOTEARS-LR and baselines across several metrics (Fig. S5b).

**Figure S5:**
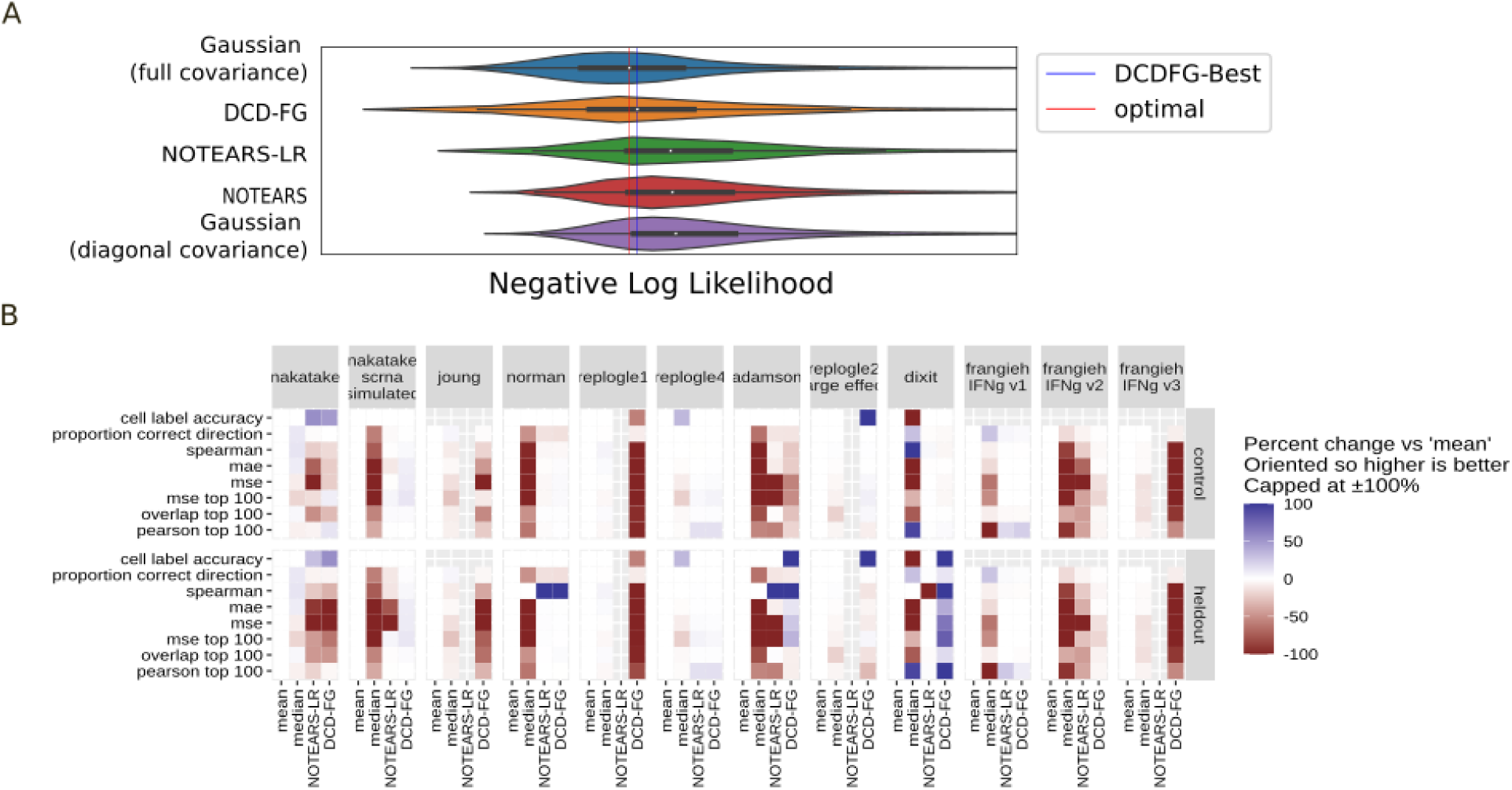
Evaluations of DCD-FG. a. Negative log likelihood on held-out perturbations in frangieh, performed using the original code for DCD-FG with Gaussian baselines added. The red line (“optimal”) indicates the best median performance across all methods. The blue line (“DCDFG-Best”) indicates the best median performance across all DCDFG-related methods. b. Benchmarks of DCD-FG and baselines (x axis) on various datasets (top margin). We use as an initial state either the control or held-out expression of upstream genes (right margin). Evaluation metrics (y axis) are as defined in Table 2. Missing data indicate that DCD-FG or NO-TEARS did not converge, or that cell type labels were not available. Performance (colorscale) is shown as percent improvement over the ‘mean’ baseline, oriented so that higher numbers always indicate better performance. Percentages are capped at ±100%.

GEARS is another expression forecasting method that has been successfully tested on held-out genetic perturbations or held-out combinations of genetic perturbations (Roohani et al., 2022). Compared to methods mentioned so far, GEARS uses a different principle to generalize to genes not perturbed in the training data: a graph neural network that enforces similar outcomes for genes with similar membership in Gene Ontology terms. Although its primary purpose is in predicting genetic interactions, GEARS has also shown robust improvement over diverse baseline methods when predicting fold change due to held-out perturbations (Roohani et al., 2022).

We examined relative performance of GEARS stratified by dataset and evaluation metric (Fig. S6). We note that three of the Perturb-seq datasets (Adamson et al., 2016; Dixit et al., 2016; Norman et al., 2019) in our collection were acquired directly from GEARS, guaranteeing identical preprocessing, although only 1,000 highly variable genes were selected (the GEARS evaluations used 6,000). GEARS performed best specifically on datasets and evaluation metrics used in the original paper (mse top 100 on adamson and norman). We note that two key factors limit this evaluation. First, GEARS is not designed for use on bulk RNA data, and PEREGGRN cannot scale to millions of cells, so for bulk data (Freimer, Nakatake) or data that were pseudo-bulked (replogle2-4, frangieh IFNg v3), no GEARS results are presented or included in aggregate performance metrics. Second, to reduce computational burden, we conducted only a single train-test split of each dataset. This could yield volatile results especially on datasets with few perturbations, such as dixit.

Finally, stratifying GeneFormer results by dataset show that it out-performed baselines on some datasets for cell label accuracy prediction, but not other metrics (Fig. S6). This corresponds to how GeneFormer was fine-tuned. We note that label prediction is not the only way of fine-tuning GeneFormer for expression forecasting, and other fine-tuning schemes may yield good performance on a wider variety of metrics. Also, datasets acquired directly from GEARS (dixit, adamson, norman) lacked raw count data, meaning that GeneFormer could not be tested on these datasets.

**Figure S6:**
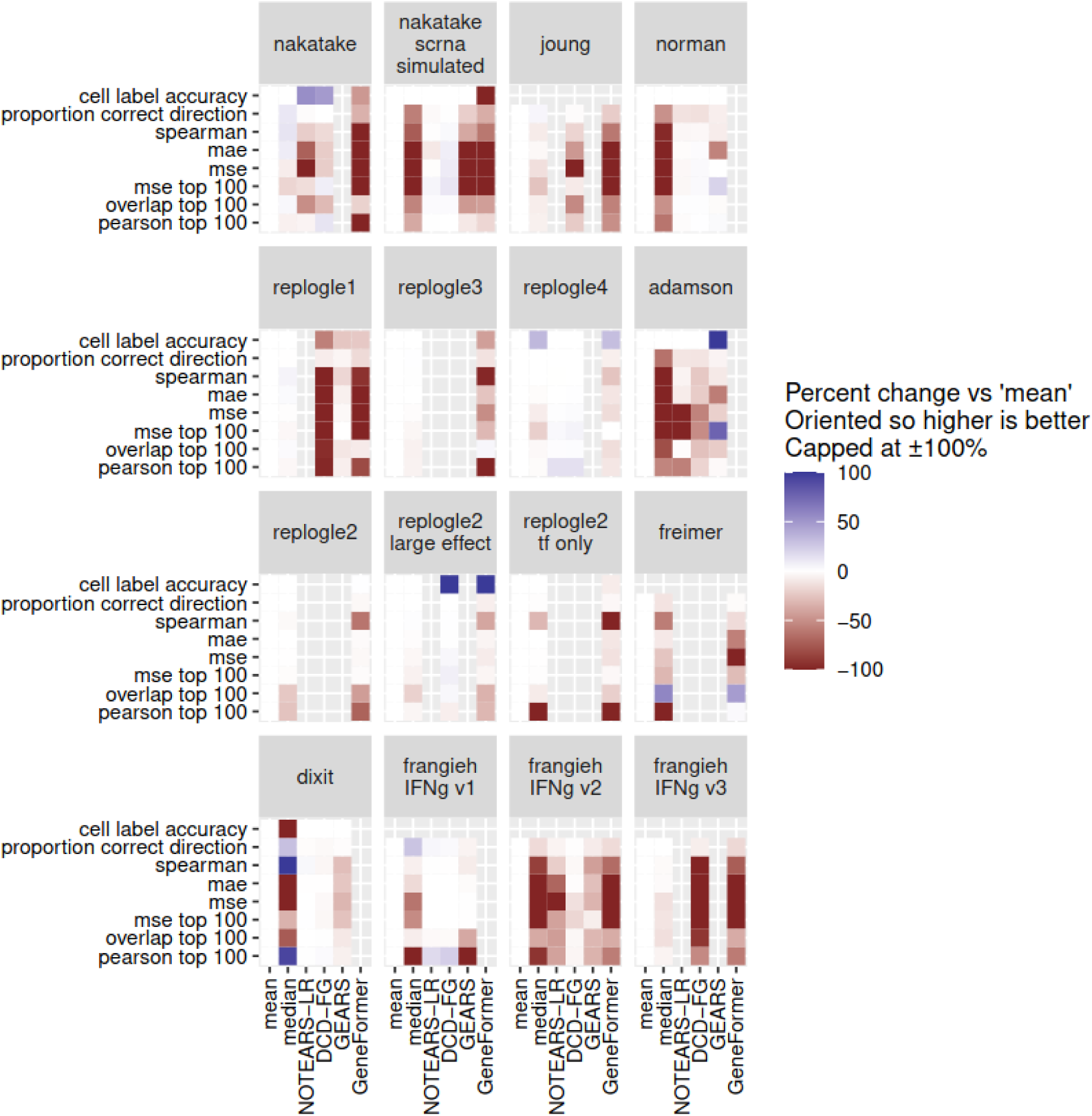
Evaluations of published expression forecasting methods (x axis) versus simple baselines. Evaluation metrics (y axis) are as defined in Table 2. Performance (colorscale) is shown as percent improvement over the ‘mean’ baseline, oriented so that higher numbers always indicate better performance. Percentages are capped at ±100%. Missing data indicate that metrics were not available, or methods did not converge, or methods cannot accept the available input data format.

## Discussion

Expression forecasting has large potential impact as a way to augment genetic screens, which are important for drug target discovery and basic science. New expression forecasting methods are being developed rapidly, and in some cases are being reported without independent validation (Amrute et al., 2022; Burdziak et al., 2023; Cui et al., 2023; Erbe et al., 2022; Hyttinen et al., 2012; Jiang et al., 2023; Kamimoto, Stringa, et al., 2023; J. Lambert et al., 2023; Osorio et al., 2022; Qiu et al., 2022; Roohani et al., 2022; Theodoris et al., 2023; Tran et al., 2022; Wang et al., 2022; Yeo et al., 2021). End users urgently need concrete evidence regarding their accuracy. To begin to answer this need, we have provided systematic expression forecasting benchmarks across a wide variety of human cell types.

Our framework enables head-to-head comparison of diverse expression forecasting methods and sources of prior knowledge. We test numerous methods using interpretable causal networks, as well as GEARS, which uses prior knowledge of functional relatedness, and GeneFormer, which is pretrained on tens of millions of single-cell transcriptomes. Most existing benchmarks focus on methods alone. In particular, the Therapeutic Data Commons genetic perturbation benchmarks include only CPA and GEARS (Velez-Arce et al., 2024), while CausalBench (Chevalley, Roohani, Mehrjou, Leskovec, & Schwab, 2022) and the affiliated GSK challenge (Chevalley, Mehrjou, Schwab, Notin, & Roohani, 2022) explicitly disallow use of prior knowledge. Although we use some of the same evaluation data as these studies, our work includes a wider range of algorithmic approaches.

One key result is that most evaluations are negative, with very simple baselines often performing at least as well as the benchmarked methods. Two very recent works (Ahlmann-Eltze et al., 2024; Wu et al., 2024) also compare pretrained transformer models, GEARS, and linear baselines on the norman, replogle3, replogle4, and other datasets. Their overall conclusions are that simpler models often outperform more complex models. Notably, these studies lack “mean” and “median” baselines, meaning their findings do not contradict our findings.

Poor perturbation prediction has important implications for competing conceptions of biological systems. Some mathematical frameworks have proven that causal models or structures can be identified even with few or no perturbations (Akutsu, Miyano, & Kuhara, 1999; Hashimoto, Gifford, & Jaakkola, 2016; Lopez et al., 2022), but these guarantees begin by assuming that all relevant causal factors have random biological variation and are perfectly observed. In other frameworks, it is not necessarily possible to predict novel perturbation outcomes (Wagner, 2001; Zhang et al., 2023), because some causal factors are unobserved, or because causal factors remain constant until reached by a perturbation. Our results indicate that the latter frameworks may be more useful conceptual models.

Another result with key implications for future studies is that expression forecasting performance evaluation has enough degrees of freedom to show many methods as the best performer, or at least better than baseline, in ways that are not robust or generalizable. Results can depend on the choice of demo dataset, the data split, and for some methods, on random initializations that cannot be fixed by the end user. Our results depended on the baselines chosen, particularly using the mean versus the median of the training data, and we expect that the mean or median of only the control samples would be an even weaker baseline. Results can also vary strongly according to the performance metric chosen. Different common-sense or previously used evaluation metrics often yielded highly discordant results. The mse, mae, and Spearman correlation tended to show baseline methods performing best -- and due to noise in the control samples, the baselines themselves can produce shockingly high correlations between observed and predicted fold change. Given that perturb-seq effects are sparse, it seems logical to predict which genes will be most affected, and then only for those genes, to predict an effect size. Focusing on the top most differentially expressed genes frequently yielded slight positive results for non-baseline methods. However, no methods consistently predicted which genes would change the most, meaning this evaluation is confined to settings where top differentially expressed genes are known. Because of these pitfalls, unbiased evaluation will require pre-specified, exactly-repeatable experiments with strong baselines and a wide variety of cellular contexts.

There are two settings where expression forecasting would be especially valuable in lieu of experiments that are impossible, expensive, or unethical. In cell types or conditions where large-scale data are not available, especially primary human cells, *in-silico* genetic screens would meet an obvious need. In cell types where many perturbations are available (Replogle et al., 2022; Subramanian et al., 2017), *in-silico* perturbation would be useful for combinatorial screening. For expression forecasting in primary cells, it would be useful to develop methods and benchmarks for using accessible cell types to assist models of less accessible cell types (Ji, Lotfollahi, Wolf, & Theis, 2021). Alternatively, natural variation in time-series or RNA velocity data could in principle be enough to reconstruct GRN’s and forecast expression, and expression forecasting based on such data has seen promising empirical results, especially when augmented with GRN structures based on motif analysis of open chromatin (Burdziak et al., 2023; Kamimoto, Stringa, et al., 2023; Pemberton-Ross et al., 2015; Qiu et al., 2022; Yeo et al., 2021). For combinatorial screening, the purpose is typically to search for cocktails of genes with specific effects such as reprogramming (Takahashi & Yamanaka, 2006). Combinatorial explosion of the search space means that expression forecasts could contribute value through scale, prioritizing a test set out of a huge number of candidate groupings (Roohani et al., 2024). Additional methods development, data generation, and benchmarking will be important in order to define and extend the limits of modern data for these specific use cases.

Enabling future extensions is crucial for bioinformatics benchmarks (Weber et al., 2019). Accordingly, a major component of this work is open source software and online documentation, with guides on not only how to repeat our results, but also how to conduct evaluations involving new methods, new datasets, new evaluation metrics, and alternative ways of splitting data that emphasize different challenges. We hope this will provide a durable resource to the expression forecasting research community.

## Methods

### Data and code availability

Our data collection is available on Zenodo at the DOI 10.5281/zenodo.8071809. Code releases used for this manuscript are in the table below. Development versions of the project infrastructure are linked from the project homepage at https://github.com/ekernf01/perturbation_benchmarking.

| Purpose | Github repository<br>(all are at ekernf01/myrepo) | DOI |
| --- | --- | --- |
| Run benchmarks | perturbation_benchmarking | 10.5281/zenodo.13862778 |
| Load and reformat networks | pereggrn_networks | 10.5281/zenodo.13862768 |
| Load and validate RNA data | pereggrn_perturbations | 10.5281/zenodo.13862772 |
| Benchmark expression forecasts | pereggrn | 10.5281/zenodo.13862784 |
| Forecast expression | ggrn | 10.5281/zenodo.13862760 |
| Extract perturbed GeneFormer embeddings | geneformer_embeddings | 10.5281/zenodo.13862756 |
| Run DCD-FG (pip-installable) | ggrn_backend2 | 10.5281/zenodo.13862764 |
| Extend original DCD-FG experiments | dcdfg | 10.5281/zenodo.13862750 |

### Simulated data (noise only)

To demonstrate that high correlation can occur between predicted and observed log fold change even when using the “mean” baseline predictor, we used the following R code, which outputs a correlation value of 0.649.

set.seed(0)

X = matrix(rnorm(10000), ncol = 100) control_indices = 1

train_indices = 1:50

test_indices = 51:100

baseline_predictor = colMeans(X[train_indices, ]) correlations = c()

for(i in test_indices){

correlations[i-50] = cor(baseline_predictor - X[control_indices, ], X[i, ] - X[control_indices, ])

}

mean(correlations)

### Simulated data (based on real networks)

During simulations, we generated data from a vector autoregressive model using the following algorithm.

- Inputs:

- by D matrix M
- by D initial states X_0_
- to perturb P
- of steps S
- Outputs:

- Simulated expression profiles of size N by D for:

- Each perturbation at the final time-point
- Each control at the final timepoint
- Each control’s initial state
- Procedure:

- For p in P:

- X_0,p_ = X_0_
- For t in 1 … S:
- X_t,p_ ^= MX^_t-1,p_
- X_t,p_[:,P] = 1
- Return X_t,p_
- Initial values

- D=811, 713, 788, or 663: the number of TF’s in the CellOracle, gtex, cellnet_human_Hg1332, or cellnet_human_Hugene networks, respectively
- M_ij_ = 1 if i regulates j in the indicated network, else 0, and M is scaled to have a maximum eigenvalue of 0.1
- X_0_ = independent random numbers each uniform on [0,1]
- P includes all genes
- S=1

### Perturbation data acquisition and preprocessing

The Nakatake data were acquired as normalized expression values by personal communication with ExAtlas maintainer Alexei Sharov (Nakatake et al., 2020). The Adamson, Dixit, and Norman datasets were obtained using GEARS. The Dixit paper describes multiple Perturb-seq experiments, but we include only the K562 TF and cell-cycle regulator experiments (Dixit et al., 2016). For all other datasets, source URL’s and citations are provided in the metadata accompanying our data collection.

Our default preprocessing included pseudo-bulk aggregation, DESeq-2 normalization, and log1p transformation. For pseudo-bulk aggregation, we summed raw counts from all cells in a given perturbation condition, reasoning that the main source of error in scRNA-seq is multinomial noise in counts and that perturbation responses in cell lines are homogeneous (Sarkar & Stephens, 2021). We also conducted exploratory analysis and clustering via scanpy (Wolf, Angerer, & Theis, 2018): we ran PCA; selected neighbors in a 100-dimensional principal subspace; and used the PCA embeddings and neighbors as input to run UMAP (McInnes, Healy, Saul, & Großberger, 2018) and the Louvain algorithm (Blondel, Guillaume, Lambiotte, & Lefebvre, 2008). Louvain clusters were omitted from freimer, frangieh, norman, and joung datasets because they did not seem attributable to perturbation effects but rather cell-cycle, donor, or depth. For knockdown and overexpression experiments, we removed samples where directly perturbed genes do not change in the expected direction. For knockout experiments, no such filter was applied, since mRNA may still be produced and may even increase due to compensatory mechanisms.

Some dataset-specific procedures differ from the above steps.

- For “nakatake”, missing values were replaced with control expression, and replication enabled testing of most perturbed genes, so we included samples only if the overexpressed gene saw a significant increase (p<0.1, t-test) or was not measured.

Nakatake scrna simulated consists of 50 simulated cells for each profile in nakatake. The expression level of cell i, gene g is Poisson with mean s_i_x_g_, where x_g_ is the normalized expression in the corresponding bulk sample and s_i_ is a size factor ensuring the total expected count is 10,000 reads per cell. No normalization is applied.
- For “freimer”, we follow the original authors and remove the outlying sample Donor_4_AAVS1_6, and we use the first ten principal components. Samples cluster by donor, so we do not evaluate methods based on Louvain cluster label predictions.
- To preprocess replogle1, we removed cells with under 2,000 UMIs or over 17,000 UMIs (99th percentile). We removed cells where over 40% of UMIs were from the 50 highest expressed genes, cells where over 20% of UMIs were from mitochondrially encoded genes, and cells where over 30% of UMIs were ribosomal protein subunits. We removed genes detected in fewer than 10 cells, unless they were overexpressed. These cutoffs are based on preliminary clustering analysis. We summed counts for each guide RNA, ran TMM normalization, and starting from the normalized pseudo-bulk expression, we removed guide RNA’s that did not increase their targets’ levels as expected.
- Preprocessing for frangieh includes three distinct versions. Version 1 follows Lopez et al. exactly to facilitate comparisons with prior work (Lopez et al., 2022). Version 2 additionally removes cells with total UMIs above the 99th percentile, with 6,000 or fewer total UMIs, with 30% or more UMIs in the top 50 expressed genes, with 10% or more UMIs from mitochondrially encoded genes, or with 20% or more UMIs from ribosomal protein subunits. Version 2 removes cells with two or more gRNAs detected, which excludes all perturbation conditions having fewer than 50 cells because the low-MOI study design only produced high cell numbers for individual gRNAs. Version 3 follows version 2, but “pseudobulk”: we added together raw counts for all cells sharing a gRNA and applied total count normalization and log1p transformation.
- Preprocessing for adamson, dixit, and norman follows GEARS (Roohani et al., 2022) to facilitate comparisons with their benchmarks; this includes only the 6,000 most variable genes in each dataset (including perturbed samples).
- We used the pseudo-bulk aggregated forms of replogle2, 3 and 4 as provided by the associated studies. We subset ‘replogle2’ as follows: Transcription factors were identified using the catalog of Lambert et al. (S. A. Lambert et al., 2018), and large effects were defined as mean_leverage_score at least 0.6.
- In joung, expression levels were summed within groups defined by TF ORF and predicted cell cycle phase, then total-count normalized.

### Gene selection

All experiments following the DCD-FG paper’s gene-selection procedure (Lopez et al., 2022). Namely, for each dataset, genes were ranked by dispersion using scanpy.pp.highly_variable_genes(…, flavor = "seurat_v3"). To select roughly N genes in a dataset with P perturbed genes, we included all genes ranked below N-P and all perturbed genes. These lists may overlap, yielding fewer than N genes. Experiments use N=10,000 by default, with the following exceptions. All experiments involving adamson, dixit, and norman include at most 6,000 genes due to upstream preprocessing (see above). All experiments involving DCD-FG use N=1000 to reduce compute time. Exact specifications for each experiment are released alongside our software (see Data and Code Availability).

### Network acquisition and preprocessing

Source URL’s, citations, and descriptions are given in the metadata accompanying our network collection, and most network acquisition is fully automated using R or python. Networks from FNTM and Humanbase were subsetted to retain only edges with posterior probability over 50%, and networks from ANANSE_0.5 or ANANSE_0.8 were subsetted to retain edges with weight over 50% or 80% respectively. GTEx coexpression networks were symmetrized: for any edge

A->B, the edge B->A was added. No other edges were added to or removed from any original source. For CellNet, starting from the documentation at http://pcahan1.github.io/cellnetr/, we downloaded the network files:

cnProc_Hg1332_062414.R

cnProc_mogene_062414.R

cnProc_Hugene_062414.R

cnProc_mouse4302_062414.R

We installed cellnetr using the instructions at https://groups.google.com/forum/#!topic/cellnet_r/pXHt2J6ZH6I. The remainder of the processing is automated.

### GRN inference methods

In initial evaluations, we use simple methods based on supervised machine learning. For each gene, a scikit-learn regressor is trained using the gene’s expression as labels and using the expression of certain TF’s as features. By default, all TF’s are included, using a manually curated list of TF’s (S. A. Lambert et al., 2018). But, if a network is used in the evaluation, then only network-adjacent TF’s are included as features. No gene is allowed to auto-regulate. Instances where gene A was perturbed directly are not used to train the model that is used to predict expression of gene A. To predict perturbation outcomes, we begin with the average expression of all controls. The corresponding transcription factor is then set to 0 (for knockout experiments) or to its observed value (for knockdown or overexpression experiments). From the resulting perturbed features, regression models are used to make predictions.

Within this same regression framework, GeneFormer is used as an alternate feature extraction method. We use GeneFormer to generate perturbed embeddings using in_silico_perturber.delete_index, in_silico_perturber.overexpress_index, and a forward pass through the model. This yields 256 features per observation. Raw counts with all genes, not normalized counts with variable genes, were passed to the GeneFormer tokenizer. When cell type labels were available, GeneFormer was fine-tuned according to the maintainers’ cell type classification examples, with a hyperparameter optimization step included.

DCD-FG and GEARS are used according to the maintainers’ documentation. GEARS was run using the “no_test” type of split unless the model could not be initialized, in which case the default type of split was used with 95% of the data used for training. NOTEARS-LR was run using DCD-FG with the LinearModuleGaussianModel. DCD-FG and NO-TEARS-LR hyperparameters were selected by reserving a simple random sample of ⅓ of the training data, then training 20 models for 600 epochs with constraint mode spectral_radius. The 20 models spanned all combinations of four latent dimensions (5, 10, 20, 50) and five LASSO penalty parameters (10, 1, 0.1, 0.01, 0.001). The combination with the best mse was selected and a model was retrained on the full training data with those parameters.

The linear embedding baseline was re-implemented in Python based on the original authors’ R code, available at https://github.com/const-ae/linear_perturbation_prediction-Paper/blob/main/benchmark/src/run_linear_pretrained_model.R. It was deployed with a ridge penalty of 0.01 and a latent dimension of 10, except on the Freimer data, where the latent dimension was reduced to 4.

### Evaluations

We attempted to test every method on all applicable datasets. Any omissions are due to RAM usage exceeding 64GB, convergence problems, runtime errors, or invalid input. Specifically, certain tree-based regression methods failed due to memory requirements exceeding 64GB. DCD-FG and NO-TEARS sometimes yielded NaN predictions across all hyperparameters. GEARS is designed for single-cell data with high sample count; we did not run it on datasets where our collection contains only bulk or pseudo-bulk expression. GeneFormer requires raw count for all genes; we did not run it on datasets where only a subset of genes were available.

For evaluation, we used the type of split marked “interventional” in the PEREGGRN documentation. Samples were grouped by perturbation, and each perturbation was allocated to the training set or the test set. Perturbations were eligible for the test set only if the perturbed gene was also measured. We used a 50-50 split, meaning 50% of perturbations (but not necessarily 50% of observations) were allocated to the test set. If fewer than 50% of perturbations were eligible for the test set, we allocated all eligible perturbations to the test set. All controls were allocated to the training set. We verified that GeneFormer was not pretrained on any of the datasets we use for evaluation.

To compute cell labeling accuracy, we trained a scikit-learn random forest classifier with 100 trees on each training dataset. The classifier was trained on the cluster labels obtained by running the Louvain algorithm (Blondel et al., 2008), which is further described in the section “Perturbation data acquisition and preprocessing”. The classifier was applied to the observed and predicted test-set expression profiles, and accuracy was measured as the fraction where classifier-assigned labels for observed and predicted expression match.

## Author contributions

E.K. initiated the study, harmonized the data, and conducted most of the benchmarking. Y.Y. benchmarked DCD-FG and performed QC on several perturbation datasets. J.W. tested code, interpreted results, and advised simulations. P.C. and A.B. funded and supervised the study. All authors edited the manuscript.

## Acknowledgements

We are grateful to Romain Lopez, Yusuf Roohani, and Christina Theodoris for generous help running DCD-FG, GEARS, and GeneFormer respectively. Thanks to Prashanthi Ravichandran and Emily Su for useful discussions and to Stephen Rosen and Dan Peng for help navigating Python packaging. Thanks to Zexi Liu for helping test the code. AB was funded by NIH grant R35GM139580. PC was funded by NIH grant R35GM124725.

## Supplemental information

Supplemental file 1: Full documentation of the Grammar of Gene Regulatory Networks

