## Supplementary material for "A systematic comparison of computational methods for expression forecasting": File S1

### GGRN: the Grammar of Gene Regulatory Networks

We describe a "grammar" capable of closely mimicking the predictive methods underlying a variety of GRN inference methods.

This grammar describes many combinations of features that are not yet implemented in our software. For clarity, these are still described in the style of software specifications. Separate web-based documentation for the GGRN python package describes a subset of options that are currently available.

#### Notation

Let $X_i(P,T)$ be RNA abundance in sample $i$, at time $T$, where genes in the set $P$ were overexpressed continuously starting at $T=0$. A natural approach is to fit $F$ such that $X_i(P,T) \approx F(X_i(P,T-\delta T))$.

#### Matching

An initial obstacle is that transcriptomics assays destroy samples: $X_i(P,T)$ and $X_i(P,T-\delta T)$ cannot both be observed. The most common work-around is to assume that samples have reached a steady state and to fit $X_i(P,T) \approx F(X_i(P,T))$; this is done by (for example) LLC, NOTEARS variants, CellOracle, Dictys, and GENIE3. If time-series data are available, an alternative is to match each sample with a counterpart at the previous time-step, which is simple with small numbers of bulk RNA samples (ScanBMA, ARMADA, DynGENIE). With single-cell RNA data, this type of matching is harder; it requires lineage tracing predictions. In PRESCIENT and scKINETICS, matching is a compromise between the predictions of the dynamic model and the observed cells nearest to those predictions. Another way to obtain paired measurements across two timepoints is RNA velocity, which is used by Dynamo. Another simple method, used by GEARS, is to match each perturbed cell with a randomly chosen control cell.

We formalize this choice with a keyword argument `matching_method` that can take the values "steady_state" (each sample is matched to itself), "closest" (each sample is matched to the closest control), "user" (matching is user-specified), "random" (each sample is matched to a random control), or "optimal_transport" (observations are matched to controls via optimal transport).

#### Regression

The matching methods described above yield a set of paired measurements $\{X_i, Y_i\}$. Every method described by this "grammar" uses supervised ML to fit $F$ such that $Y_i \approx F(X_i)$. Different families are assumed: CellOracle, ScanBMA, Dictys, SCKINETICS, ARMADA, LLC, NOTEARS, and NOTEARS-LR use linear regression; DCD-FG uses multilayer perceptrons; NOBEARS uses polynomial regression; GENIE3 uses random forests; Dynamo uses kernel ridge regression; and PRESCIENT and RNAForecaster use neural networks. Sometimes these are not trained directly on expression levels; they may also involve projection into lower-dimensional latent spaces. Low-dimensional structure will be discussed shortly.

We formalize the choice of regression method with a keyword argument `regression_method` that can take values such as `RidgeCV` or `LassoCV`. More options may be added by specific software implementations and should be clearly documented alongside the software.

#### Feature extraction and TF activity

Many transcription factors are post-transcriptionally regulated, and the mRNA level may not be the best proxy for TF activity. Thus, although it is still common to use mRNA levels, there are other possible choices of features used in training the model $F$. GeneFormer uses a 256-dimensional latent space representation for each cell. PRESCIENT uses a 30- or 50-dimensional principal subspace embedding. ARMADA augments each log-normalized expression profile $X$ with additional coordinates $A$ representing TF activity. $A$ is estimated by fitting $X_{ij} \approx \sum_m N_{jm} A_{mi}$, where $N_{jm}$ counts how many times motif $m$ occurs near the promoter of gene $j$. $N_{jm}$ is constructed from promoter annotations, motif databases, and DNA sequence; it does not depend on expression data except through possibly-novel promoter annotations.

For mRNA levels, the effect of genetic perturbation can be directly measured (although knockouts may have nonzero mRNA levels despite a complete loss of function). For more complex feature extraction, each model must specify how genetic perturbations affect the features. GeneFormer uses a tokenized representation of each cell which orders genes according to a measure of expression anomaly over baseline; for perturbations, GeneFormer places overexpressed genes first and removes deleted genes, then generates embeddings. GEARS learns gene embeddings during training, using latent space arithmetic to alter embeddings of perturbed cells. PRESCIENT projects into the principal subspace after altering z-scores for perturbed genes. ARMADA does not simulate perturbations, but ARMADA has uniquely interpretable motif activity scores and perturbations could be directly applied to individual motif activities.

We formalize these choices with a keyword argument `feature_extraction` that can take the values `mRNA` or `GeneFormer`. More options may be added by specific software implementations and should be clearly documented alongside the software.

#### Low dimensional structure

Some methods' estimates of $F$ are heavily constrained by low-rank structure. Furthermore, different methods provide different frameworks to describe how low-dimensional projections inter-operate with dynamic models. Let $Q(X)$ project $X$ into a $D$-dimensional space (via PCA unless otherwise noted). Let $R$ be a right inverse of $Q$ (this means $Q(R(x)) = x$ for all $x$). Let $G$ be a model for the progression of the cell's state over one time-step. $G$ will sometimes be applied in expression space and sometimes in lower dimensional space. We summarize some different approaches in terms of the order in which $R$, $G$, and $Q$ are applied.

- CellOracle models dynamics in the original space, but at the final step requires projection of all output into a low-dimensional space for biological interpretation of predictions. CellOracle is not intended to reconstruct full gene expression profiles, but two sensible options would be to reconstruct expression from the final results in the PCA subspace, which GGRN would represent as $F(X) = R(Q(G(X)))$, or to never project into the PCA subspace, which GGRN would represent as $F(X) = G(X)$. Here, $G$ makes predictions using the Bayesian ridge regression models trained by CellOracle.
- PRESCIENT models dynamics after projection into a 30- or 50-dimensional principal subspace, and to obtain per-gene expression predictions, it would be necessary to project back into higher-dimensional space such that $F(X) = R(G(Q(X)))$.
- ARMADA assumes $F(X) = R(G(Q(X)))$, but the "encoder" $Q$ is not learned via PCA or backprop. Rather, it is fixed *a priori* based on motif analysis; in the notation above, $Q(X) = N^{\dagger}X$ where $N^{\dagger}$ is the Moore-Penrose pseudoinverse of the motif-by-promoter count matrix $N$.
- RNAForecaster's neural ODE specification includes a hidden layer of user-defined size. The default size is larger than the input dimension, but if users provide smaller values, this could be used as a low-dimensional constraint on predicted expression.
- DCD-FG and NOTEARS-LR each jointly learn an "encoder" $Q$ and a "decoder" $D$ such that $F(X) = D(Q(X))$. The decoder effectively plays the role of both $R$ (projecting a low-dimensional vector back to the original data dimension) and $G$ (modeling dynamics). For NOTEARS-LR, $D$ and $Q$ are linear, and for DCD-FG, $D$ is linear but $Q$ consists of $K$ separate multilayer perceptrons, where $K$ is the latent dimension. NOTEARS-LR and DCD-FG do not use PCA; they learn the encoder and the decoder in a supervised way by backpropagating errors.

This variety can be mostly captured by two key questions. First, can the assumed dynamics be fully described by an action on the low-dimensional space? For CellOracle, no; for ARMADA, PRESCIENT, and DCD-FG, yes. Second, how are the low-dimensional projections learned? For ARMADA, they are fixed; for CellOracle and PRESCIENT, they are learned by PCA; for DCD-FG, they are learned by back-propagating errors. We formalize these options with keyword arguments:

- `low_dimensional_structure` can take values:
  - `none` , meaning $F(X) = G(X)$ ,
  - `dynamics` , meaning $F(X) = R(G(Q(X)))$,
  - `postprocessing` , meaning $F(X) = R(Q(G(X)))$.
- `low_dimensional_training` can take values `user` ($Q$ is the pseudo-inverse of a user-provided matrix, as in ARMADA), `pca` (Q is learned by running PCA on the training data, as in PRESCIENT or CellOracle), or `supervised` ($Q$ and $R$ are learned from the input-output pairs, as in DCD-FG).
- `low_dimensional_value` can be a positive integer giving the latent dimension or an entire matrix if `low_dimensional_training` is `user` .

#### Sparse structure

Estimates of $F$ are also heavily affected by assumptions about sparsity. DCD-FG and NOTEARS variants use an L1 penalty so that each coordinate of $F(X)$ (or $Q(X)$) is a function of just a few coordinates of $X$. ScanBMA uses Bayesian model scoring tools toward the same end, selecting a model with high BIC and averaging its predictions with nearby models above a certain performance threshold. CellOracle and SCENIC use ML methods that yield dense solutions, but they each include a step to prune weak coefficients and re-fit each regression with fewer inputs. LLC uses an L1 penalty to induce sparsity for some applications. PRESCIENT and Dynamo do not assume any sparsity.

Sparsity can arise from certain choices of `regression_method` described above, such as `LassoCV` . GGRN also includes keyword args:

- `pruning_strategy` can be `"none"` or `"prune_and_refit"` .
- `pruning_parameter` is a floating point number.

If `pruning_strategy` is `"prune_and_refit"` , then all coefficients with magnitude less than `pruning_parameter` are fixed to zero, and the models are re-fit.

#### Feature selection

Some models (CellOracle, Dictys) use only transcription factors as features when learning $F$. Others (e.g. DCD-FG) use all genes. We include a keyword arg `eligible_regulators` which accepts a list of all features to use when learning $F$.

#### Known or suspected interactions

The specific pattern of connections in the network can be informed by prior knowledge about regulatory relationships -- most often, motif analysis. CellOracle, Dictys, ARMADA, and SCENIC+ allow regulation of gene $j$ by regulator $k$ only if a matching motif is found near the promoter of $j$ or in a paired enhancer. Earlier iterations of ScanBMA used a similar hard *a priori* threshold, while later ScanBMA papers include the same information but as an informative prior rather than a hard threshold, meaning edges outside the prior-knowledge network may be added if there is sufficient evidence in the training data.

We formalize these alternatives with a keyword argument `network` that accepts a container of user-input TF-target ordered pairs (in our software, this is always a memory-efficient LightNetwork object). The edges may be weighted and they may be cell type-specific. We also include an argument `network_prior` that can take the values `ignore`, meaning `network` will not be used, or `restrictive`, meaning gene $j$ is predicted by regulator $k$ only if `network` contains that TF-target pair. In the future intermediate options like ScanBMA could be added.

#### Autoregulation

Some methods allow autoregulation, while others (e.g. DCD-FG, CellOracle) must exclude autoregulation to obtain non-trivial solutions. To represent this, we include a keyword arg `predict_self` taking the values True or False.

#### Cell-type specificity

Many perturbation prediction methods use the same model across each dataset. These tend to be models that were demonstrated on fairly homogenous systems like yeast (ScanBMA), hematopoeisis (PRESCIENT, Dynamo) or THP-1 or HUVEC cells (ARMADA). However, CellOracle and Dictys can be deployed on more heterogeneous systems (e.g. whole zebrafish embryos), and they opt to train a separate $F$ on each of many cell types or developmental stages. For CellOracle, the regression, pruning, and refitting are done separately within each cell type. An intermediate option would be to fit cell type-specific models, but shrink each model towards shared structures or shared effect sizes.

We formalize these alternatives with a keyword argument `cell_type_sharing_strategy` that can take the values `identical` (one model used throughout) or `distinct` (one model per cell type). Additional options may be added in the future to describe specific strategies for partial sharing of effects or structure across cell types.

#### Predictive timescale

To predict the effect of perturbing gene $j$ to level $z$ (e.g. $z = 0$ for a knockout), an obvious choice is to start with control expression $X$, set $X_j \leftarrow z$, and then set $X \leftarrow F(X)$. For models like Dynamo, PRESCIENT, ARMADA, and ScanBMA that are coupled to an explicit time-scale, an entire trajectory can be computed by iterating this process. For steady-state models, the amount of time simulated by a single iteration is unclear. In DCD-FG, typically a single iteration is used. CellOracle leaves this to the user, recommending 3 iterations. Dictys proceeds until a steady state is reached.

We formalize this with a keyword argument `prediction_timescale` that accepts a list of positive numbers specifying the number of steps to take. Preferably this should be specified on the same scale as the time-point labels in the training data.

#### Perturbation persistence

In PRESCIENT, $X_j = z$ is done once, and only $X = F(X)$ is iterated, corresponding to a transient perturbation (e.g. inducible knockdown). In CellOracle, each iteration includes both $X_j = z$ and $X = F(X)$, corresponding to a persistent perturbation (e.g. knockout). In Dynamo, perturbations also persist; they are incorporated into the entire vector field of derivatives, and a path is selected to minimize the "action", a measure of discordance between the curvature of the path and the predicted vector field of post-perturbation derivatives. LLC also assumes persistent perturbations.

We formalize this with a keyword argument `do_perturbations_persist` that is Boolean.

#### Interpretation of randomness

RNA-seq, and especially single-cell RNA-seq, produce data that are noisy. Transcription itself is also stochastic. Most of the methods described here include random elements, which could be interpreted as biological reality or measurement error. PRESCIENT, Dictys, and DCD-FG each propagate noise from gene to gene in their generative models, meaning randomness is interpreted as biological reality. Dictys includes boh biological stochasticity via a stochastic differential equation and a binomial model for measurement error. Due to the difficulty of causal inference on latent variables, GGRN currently neglects measurement error and biological stochasticity.

#### Summary

GGRN can describe methods by:

- `matching_method` : "steady_state", "closest", "user", "random", "optimal_transport"
- `do_perturbations_persist` : boolean
- `prediction_timescale` : list of positive numbers
- `regression_method` : A method from sklearn, e.g. "KernelRidge"
- `eligible_regulators` : A list of all genes to use as features; often a list of all TFs
- `feature_extraction` : "mRNA" or "GeneFormer", possibly with additional options in the future
- `predict_self` : False to disallow autoregulation, True otherwise
- `low_dimensional_structure` : "none", "dynamics", or "postprocessing"
- `low_dimensional_training` : "user", "svd", "supervised"
- `pruning_strategy` : "none", "prune_and_refit"
- `network` : An edge list containing regulators, targets, weights (optional), and cell types (optional)
- `network_prior` : "ignore" or "restrictive"

- `cell_type_sharing_strategy` : "identical", "distinct", possibly with additional options in the future

#### Limitations

GGRN cannot fully represent the distinctive features of most methods. Some specific examples where GGRN falls short:

- This document outlines more features than are yet implemented in our software. Separate documentation describes a subset of options that are currently available.
- GGRN is limited to discrete-time by the initial matching step, and can only approximate continuous-time methods (PRESCIENT, Dictys, Dynamo). For a detailed explanation of connections between discrete-time linear models and continuous-time nonlinear models consult Oates & Mukherjee 2012.
- GGRN cannot specify the exact network architectures used in DCD-FG's factor graph, PRESCIENT's potential function, or GEARS.
- GGRN cannot capture how PRESCIENT and scKINETICS alternate between matching and training. These methods do not match $X_i(T - \delta T)$ to $X_j(T)$. Instead they match $F(X_i(T - \delta T))$ to $X_j(T)$, jointly optimizing over both $F$ and the matching method's output.
- GGRN lacks any description of proliferation and apoptosis. PRESCIENT includes features for modeling proliferation and apoptosis, which are essential to its performance in benchmarks based on clonal barcoding.
- GGRN does not describe any data preprocessing, rather assuming log normalized gene expression is given as input.
- GGRN does not describe any motif analysis or chromatin data analysis; systematizing that domain would be a separate endeavor. To highlight specific important features not included in GGRN: CellOracle, SCENIC+, and Dictys include bespoke pipelines for motif analysis and data-driven pairing of enhancers with promoters. ARMADA emphasizes promoter annotation, models of motif evolution, and motif positioning relative to the promoter.
- GGRN cannot specify the details of foundation models pre-trained on large collections of transcriptome data; systematizing that domain would be a separate endeavor.
- GGRN does not describe acyclic penalties such as those used by DCD-FG and related causal inference methods.
